# Validation of a Commercially Available Non-Woven Fabric Electrode for Mouse Electroretinography

**DOI:** 10.64898/2026.10.07.757191

**Authors:** Yuta Ishizuka, Hiroyuki Kamao, Junko Matsuda

**Affiliations:** Department of Pathophysiology and Metabolism, Kawasaki Medical School, Okayama, Japan; Department of Ophthalmology, Kawasaki Medical School, Okayama, Japan

## Abstract

**Purpose:** To evaluate the performance and usability of a commercially available non-woven fabric electrode as a cost-effective and user-friendly alternative to standard corneal contact lens electrodes for recording murine electroretinograms.

**Methods:** Using a paired design, full-field electroretinograms were recorded in the same mice using the commercially available non-woven fabric electrode and a conventional contact lens electrode. Key parameters evaluated included (1) stability of baseline noise, (2) dark-adapted intensity–response curves (0.01–100 cd·s·m⁻²), (3) light-adapted intensity–response curves (3–100 cd·s·m⁻²), (4) 20-Hz and 30-Hz flicker responses, (5) extended recordings targeting slow c-wave kinetics under dark-adapted conditions, and (6) inter-eye consistency via bilateral simultaneous recordings.

**Results:** Eliminating the need for viscous lubricants, the non-woven fabric electrode simplified placement on the cornea, alignment, and re-application. It yielded a-wave and b-wave amplitudes and peak times comparable or superior to those obtained with standard contact lens electrodes. Notably, during prolonged recordings, the non-woven fabric electrode yielded significantly larger c-wave amplitudes while maintaining equivalent baseline noise stability. Bilateral simultaneous recordings demonstrated high inter-eye correlation and low variability of measurements between contralateral eyes.

**Conclusions:** The non-woven fabric electrode provides high-quality, reproducible electroretinographic recordings comparable to (and for c-waves superior to) those obtained using standard corneal contact lens electrodes. This method facilitates standardized data acquisition and improves inter-laboratory reproducibility of murine electroretinography.

**Translational Relevance:** By reducing financial and technical barriers, this non-woven fabric electrode offers an alternative to specialized recording setups, improving experimental throughput and efficiency in ophthalmic research.

## Introduction

First described by Holmgren in 1866,^1^ electroretinography (ERG) has evolved into an indispensable non-invasive method for evaluation of retinal function. Since the International Society for Clinical Electrophysiology of Vision (ISCEV) published its first standardized protocol to improve the comparability of inter-laboratory data,^2^ full-field ERG standards have been regularly updated.^3–5^ Building upon these human clinical standards, standardized protocols have also been established and used extensively in rodent models, particularly mice and rats, to evaluate physiological retinal function and investigate pathological mechanisms in retinal diseases.^6–10^

Traditionally, contact lens electrodes have been regarded as the *de facto* standard for rodent ERG because of their high signal amplitude and reproducibility of recording.^6,9,11,12^ Automated all-in-one platforms, such as the Celeris system, are also commercially available.^10,13^ While these dedicated platforms provide standardized data, integrated systems require high initial equipment costs. Furthermore, application of contact lens electrodes to the eyes of small animals presents technical challenges, requiring precise alignment and posing risks of corneal desiccation or altered intraocular pressure as a result of excessive mechanical pressure.^8,14,15^

Soft cotton electrodes have been explored over many years in an effort to circumvent the technical drawbacks of contact lens electrodes.^12,14,16^ While soft cotton electrodes yield robust responses with high signal amplitudes and offer superior corneal compatibility, the traditional versions were manually fabricated by encasing saline-soaked cotton within syringe barrels or glass tubes to maintain electrolyte hydration or by hand-rolling fine cotton-wicks around metal lead wires. Therefore, these manually crafted electrodes lack structural consistency, require frequent re-wetting during testing, and are difficult to sterilize or reuse, thereby hindering inter-laboratory standardization. Furthermore, these structural inconsistencies can adversely affect the key biophysical properties of the electrode, such as impedance, the noise floor, and baseline potential drift, which may compromise signal stability, particularly during multi-second continuous recordings.^11^

In an effort to overcome both the drawbacks associated with corneal contact lens electrodes and the structural variability of handmade cotton electrodes, we repurposed a commercial auditory canal electrode containing a silver-based non-woven fabric pad (CE electrode, DS-101-300UO; Unique Medical, Tokyo, Japan) as a mouse corneal active electrode. This pre-fabricated electrode retains saline through capillary action and maintains corneal hydration without requiring viscous lubricants, while allowing easy alignment and reuse after re-sterilization. In this study, we performed paired comparisons of this commercial non-woven fabric electrode and a standard corneal contact lens electrode in the same mice. We evaluated full-field ERG responses across dark-adapted, light-adapted, and flicker protocols, including baseline noise levels, multi-second c-waves, and inter-eye correlations, to validate its reliability and reproducibility when performing visual electrophysiology tests in mice.

## Methods

### Experimental Animals

Seven adult C57BL/6J mice (4 male, 3 female; age 9–14 weeks) were used in the study. They were housed in a temperature-controlled facility (23 ± 2°C) under a 12-h light/dark cycle (lights on at 08:00) with access to food and water *ad libitum*. All animal handling and experimental procedures were approved by the Institutional Animal Care and Use Committee of Kawasaki Medical School (approval number 25-011) and performed in compliance with the Institutional Guidelines for Animal Experiments, ARRIVE guidelines, and ARVO Statement for the Use of Animals in Ophthalmic and Vision Research.

### Animal Handling and Anesthesia Protocol for Electroretinography

The mice were transferred to the ERG recording room and dark-adapted overnight (for >16 h). All subsequent procedures were performed under dim red illumination. Surgical anesthesia was induced via a subcutaneous injection of a medetomidine-midazolam-butorphanol mixture containing medetomidine hydrochloride (0.38 mg/kg), midazolam (4.8 mg/kg), and butorphanol tartrate (5.8 mg/kg) at a volume of 10 mL/kg. Throughout the experimental session, each mouse was maintained on a homeothermic heating pad (ThermoStar Homeothermic Control System; RWD Life Science, Shenzhen, China) set to 40°C to prevent hypothermia. Before placement of the electrodes, the pupils were fully dilated with topical 0.5% tropicamide/0.5% phenylephrine hydrochloride eye drops (Mydrin-P; Santen Pharmaceutical, Osaka, Japan). Local corneal anesthesia was achieved using 0.4% oxybuprocaine hydrochloride eye drops (Benoxil; Santen Pharmaceutical).

### Recording of ERG responses

Full-field ERG responses were recorded simultaneously from both eyes using a data acquisition system (PowerLab; ADInstruments, Sydney, NSW, Australia) coupled with two BioAmp amplifiers (ADInstruments) connected in parallel for two-channel acquisition and a small-animal Ganzfeld stimulator driven by an LED light source (Rat/Mouse Ganzfeld and LS-100; MAYO, Aichi, Japan). A stainless steel needle reference electrode was inserted subcutaneously in the snout, and a ground needle electrode was placed subcutaneously in the midline of the head between the ears. For active recording, either a conventional corneal contact lens electrode (MAYO) or a CE electrode was placed on the corneal surface, with sterile normal saline (Otsuka Pharmaceutical, Tokyo, Japan) applied topically to keep the cornea hydrated and ensure optimal electrical contact. The conventional contact lens electrode was manually positioned directly on the cornea. Representative photographs of the components of the CE electrode, a structural comparison of the CE electrode with the conventional contact lens electrode, and the positioning of the CE electrode on the mouse eye are shown in Figure 1. The CE electrode was initially positioned using movable simple manipulators placed adjacent to both eyes (Figure 1C); the electrode was then brought into contact with the cornea via the saline droplet, followed by removal of the excess saline to achieve stable fitting (Figure 1D). Amplifier bandpass filters were set to 0.1–500 Hz to record raw ERG signals with minimal filtering and prevent attenuation of slow response components, such as the c-wave, and oscillatory potentials were not eliminated for analysis. Signals were digitized at a sampling frequency of 2 kHz (with a time resolution of 0.5 ms).

**Figure 1.**
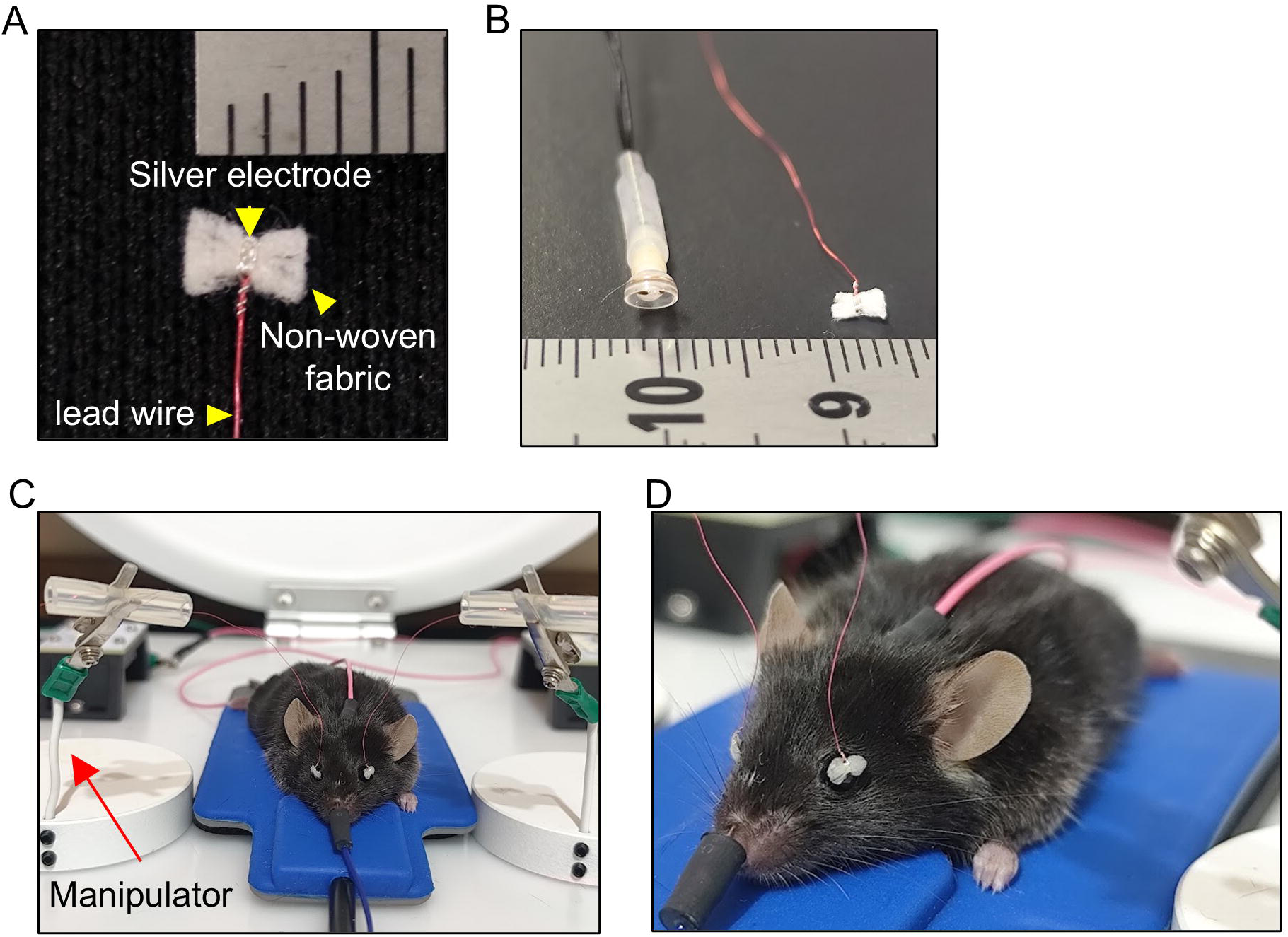
(A) Photograph showing the components of the CE electrode (silver electrode, non-woven fabric, and lead wire) alongside a millimeter ruler. (B) Comparison of size between a conventional contact lens electrode (left) and a CE electrode (right) alongside a millimeter ruler. (C) Representative photograph showing the setup with a CE electrode placed on each eye of a mouse. (D) Close-up lateral view showing the CE electrode fitted correctly onto the mouse cornea.

Dark-adapted (DA, scotopic) ERG series were evoked by brief flash stimuli ranging from 0.01 to 100 cd·s·m⁻² (DA 0.01 to DA 100). For dark-adapted stimulus intensities up to 30 cd·s·m⁻² (DA 30), three consecutive responses were recorded and averaged for each condition. Single-flash responses without averaging were recorded at 100 cd·s·m⁻² (DA 100) to prevent signal attenuation resulting from repeated intense photic stimulation. For evaluation of the background noise level for each electrode, the root mean square amplitude was calculated from the pre-stimulus baseline period (1.0 s before the flash) of the lowest dark-adapted intensity responses (DA 0.01).

Following the scotopic measurements, the mice were light-adapted for >5 min to a steady, rod-saturating background field of 31.6 cd·m⁻². Light-adapted (LA, photopic) single-flash ERGs were recorded at stimulus intensities ranging from 3 to 100 cd·s·m⁻² (LA 3 to LA 100) against the continuous background field. For each stimulus intensity, signals from 32 repeated flashes were averaged to obtain the final response waveform. For waveform analysis, the average potential recorded during the 20-ms pre-stimulus period (−20 to 0 ms) was defined as the baseline.^5^ The a-wave was identified as the initial negative deflection reaching its trough within 5 ms to 35 ms post-flash, and its amplitude was measured from the baseline to the trough (in μV). The b-wave was identified as the positive peak occurring after the a-wave, and its amplitude was measured from the a-wave trough to the b-wave peak (in μV). Peak times (also called implicit times) for both a-waves and b-waves were defined as the times elapsed from the stimulus flash to their respective troughs and peaks. Photopic flicker ERGs were subsequently elicited by 20-Hz and 30-Hz flicker stimuli at 3 cd·s·m⁻² (LA 3) under the same 31.6 cd·m⁻² background field. Ten consecutive series, each comprising 20 flash stimuli, were recorded for each frequency condition. To avoid transient onset/offset artifacts, response waveforms were evaluated using the middle 10 flashes of each series, excluding the first and last 5 flashes. Data obtained from the left and right eyes were averaged for each animal and used as the representative value for statistical analysis.

Following completion of the recording session, atipamezole hydrochloride (4.0 mg/kg, administered subcutaneously at a volume of 10 mL/kg) was injected to reverse the anesthesia induced by medetomidine-midazolam-butorphanol. Each animal was maintained on a warming plate (40°C) until complete recovery from anesthesia and received topical 0.1% sodium hyaluronate solution, prepared by dissolving sodium hyaluronate powder (FUJIFILM Wako Pure Chemical Corporation, Osaka, Japan) in normal saline, to prevent corneal desiccation.

### Benchtop Electrical Impedance Measurement

The electrical impedance of conventional contact lens electrodes and CE electrodes (*n* = 2 each: unit 1 and unit 2) was evaluated using a chemical impedance analyzer (IM3590, Hioki, Nagano, Japan). The electrodes were immersed in normal saline at room temperature, and impedance spectra were recorded across a frequency range from 1 Hz to 1 kHz. Five consecutive measurements (technical replicates) were performed for each unit to assess the reproducibility of measurements and consistency between the units.

### Statistical Analysis

The sample size was determined by an a priori power analysis using EZR software.^17^ Considering the paired design in which two electrodes were compared within the same individuals,^12^ we calculated that a minimum of six mice would be required to achieve over 80% power at a significance level of 0.05 for detection of an estimated mean difference of 150 µV in a-wave amplitude with a standard deviation of 100 µV. This sample size aligns with established protocols recommending at least five animals for evaluations of murine ERG.^6^ Therefore, seven mice were included to account for potential loss of data while ensuring robust statistical power and adhering to the 3Rs principles of animal research.

Intensity–response data for ERG amplitudes were fitted with non-linear regression using a sigmoidal Naka–Rushton function. A global fitting analysis was performed to determine whether the overall intensity–response curves differed significantly between the contact lens electrode group and the CE electrode group, comparing a single shared curve against separate curves for each group using the extra sum-of-squares *F*-test. The Wilcoxon signed-rank test was used for pair-wise comparisons between the two electrodes at individual stimulus intensities. Linear regression was performed on simultaneously recorded bilateral ERG amplitudes to determine regression equations and coefficients of determination (R²) for assessment of inter-eye consistency and the reliability of recording. All values are expressed as the mean ± standard error of the mean (SEM). All data processing and statistical analyses were performed using GraphPad Prism version 7.05 (GraphPad Software Inc., San Diego, CA, USA). A *P*-value of < 0.05 was considered statistically significant.

## Results

### Analysis of Baseline Noise

We quantified the electrical baseline stability of both recording systems (Figure 2A and 2B) before evaluating the ERG responses. There was no significant difference in root mean square noise (calculated from the 1.0-s pre-stimulus) between the contact lens electrode and the CE electrode (12.08 ± 1.20 μV vs. 14.79 ± 1.81 μV, *P* = 0.2188; Figure 2C). Both electrodes maintained consistently low baseline noise levels, demonstrating that the CE electrode provides electrical contact stability similar to that of the conventional contact lens electrode.

**Figure 2.**
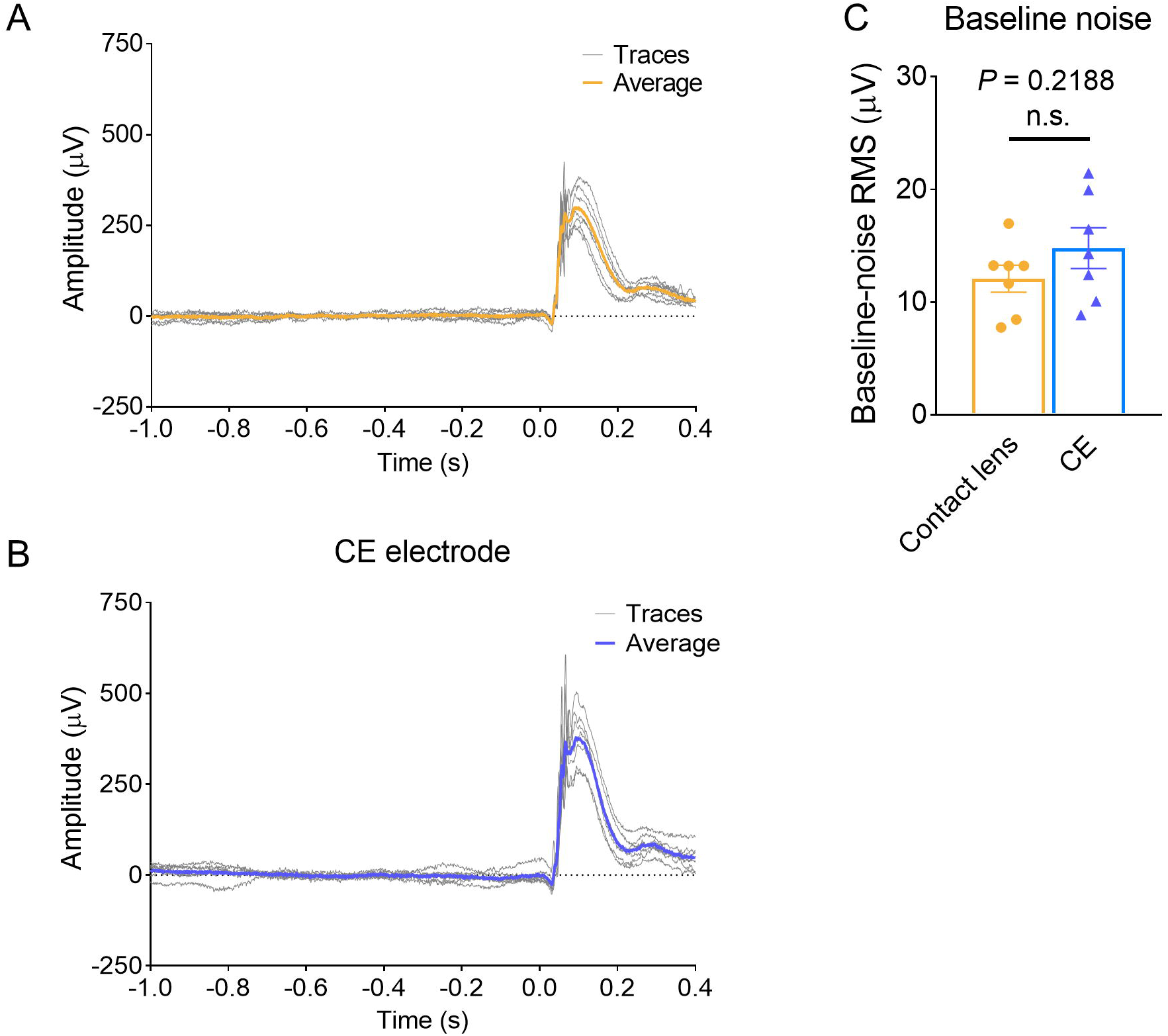
Individual electroretinographic traces from all subjects and their corresponding averaged waveforms, along with a quantitative comparison of baseline noise levels between conventional contact lens electrodes and CE electrodes. Individual electroretinographic traces (gray lines) spanning from −1.0 s to +0.4 s relative to stimulus onset and their corresponding average waveforms (colored lines) recorded with a contact lens electrode (yellow, A) and a CE electrode (blue, B) (*n* = 7). (C) Quantitative comparison of RMS baseline noise amplitude measured before stimulus onset (mean ± standard error of the mean, *n* = 7). Statistical significance was determined using the Wilcoxon matched-pairs signed-rank test (*P* = 0.2188; n.s., not statistically significant). RMS, root mean square

### Dark-Adapted and Light-Adapted ERG Waveforms

Next, we evaluated averaged dark-adapted and light-adapted single-flash ERG waveform series recorded with the conventional contact lens electrode and the CE electrode across a range of flash intensities (Figure 3). Under dark-adapted conditions (Figure 3A and 3B; mean waveforms with shaded SEM regions are shown in Supplementary Figure S1), increasing the stimulus intensity from 0.01 to 100 cd·s·m⁻² elicited characteristic intensity-dependent waveform changes in both electrode groups; low-luminance flashes produced broad b-waves, whereas higher-luminance flashes (>0.3 cd·s·m⁻²) induced prominent negative a-wave deflections followed by robust b-waves. Similarly, under light-adapted conditions (3–100 cd·s·m⁻², Figure 3C and 3D; mean waveforms with the SEMs depicted as shaded regions are shown in Supplementary Figure S2), photopic responses showed consistent luminance-dependent increases in peak amplitudes for both recording configurations. Notably, while the overall waveform morphology, peak times, and signal-to-noise dynamics recorded by the CE electrode closely paralleled those obtained using the contact lens electrode across all tested conditions, the CE electrode consistently showed slightly higher response amplitudes. Quantitative statistical comparisons of a-wave and b-wave amplitudes as well as peak times are presented in Figure 4.

**Figure 3.**
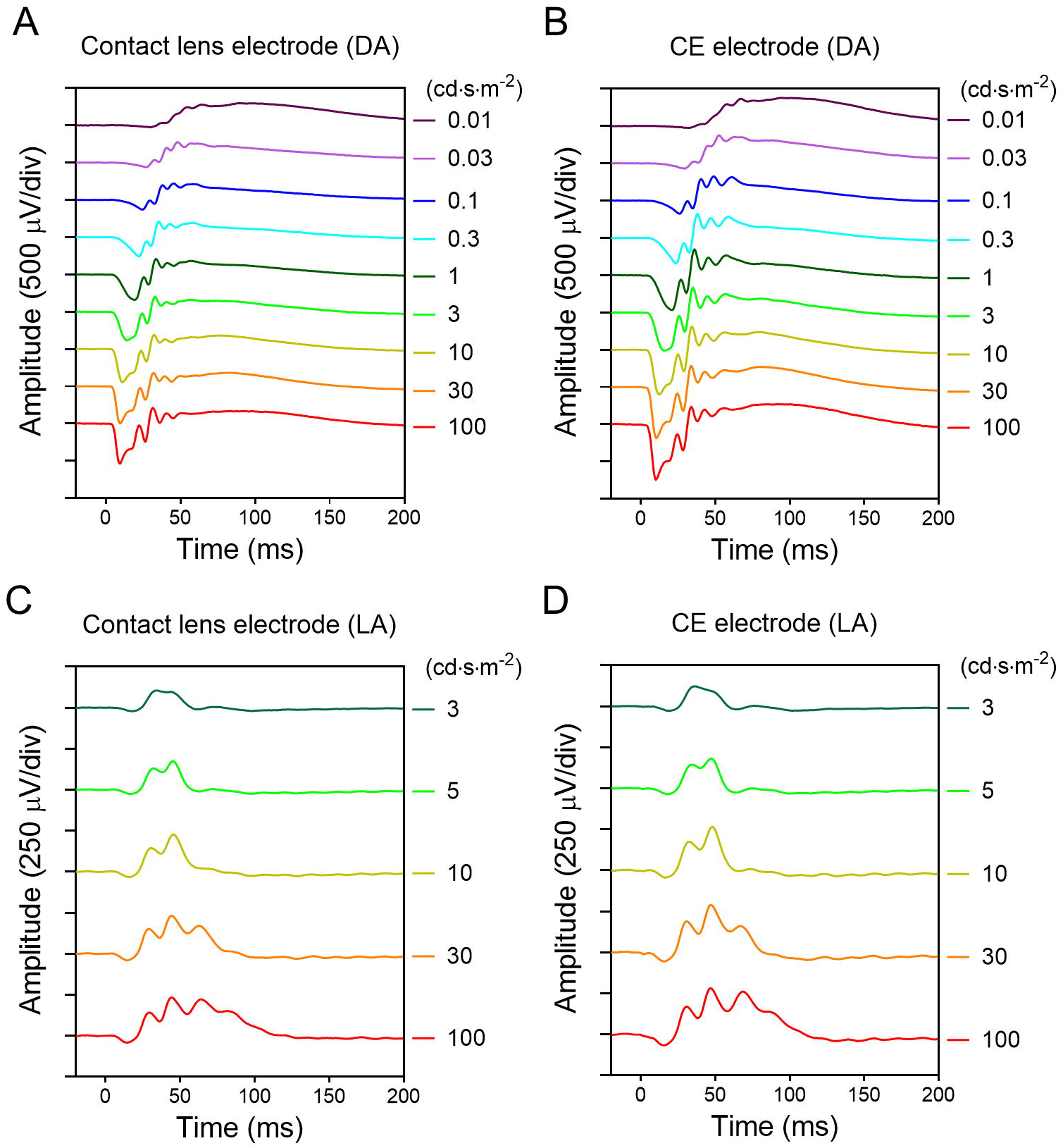
Intensity–response series of dark-adapted (DA) and light-adapted (LA) electroretinograms recorded using contact lens and CE electrodes. A representative dark-adapted electroretinographic trace series recorded with contact lens electrodes (A) and CE electrodes (B) across a flash intensity range of 0.01 to 100 cd·s·m⁻². Traces are stacked vertically in ascending order of flash intensity. Representative light-adapted electroretinographic trace series recorded on a constant background light with contact lens (C) and CE (D) electrodes across a flash intensity range of 3 to 100 cd·s·m⁻².

**Figure 4.**
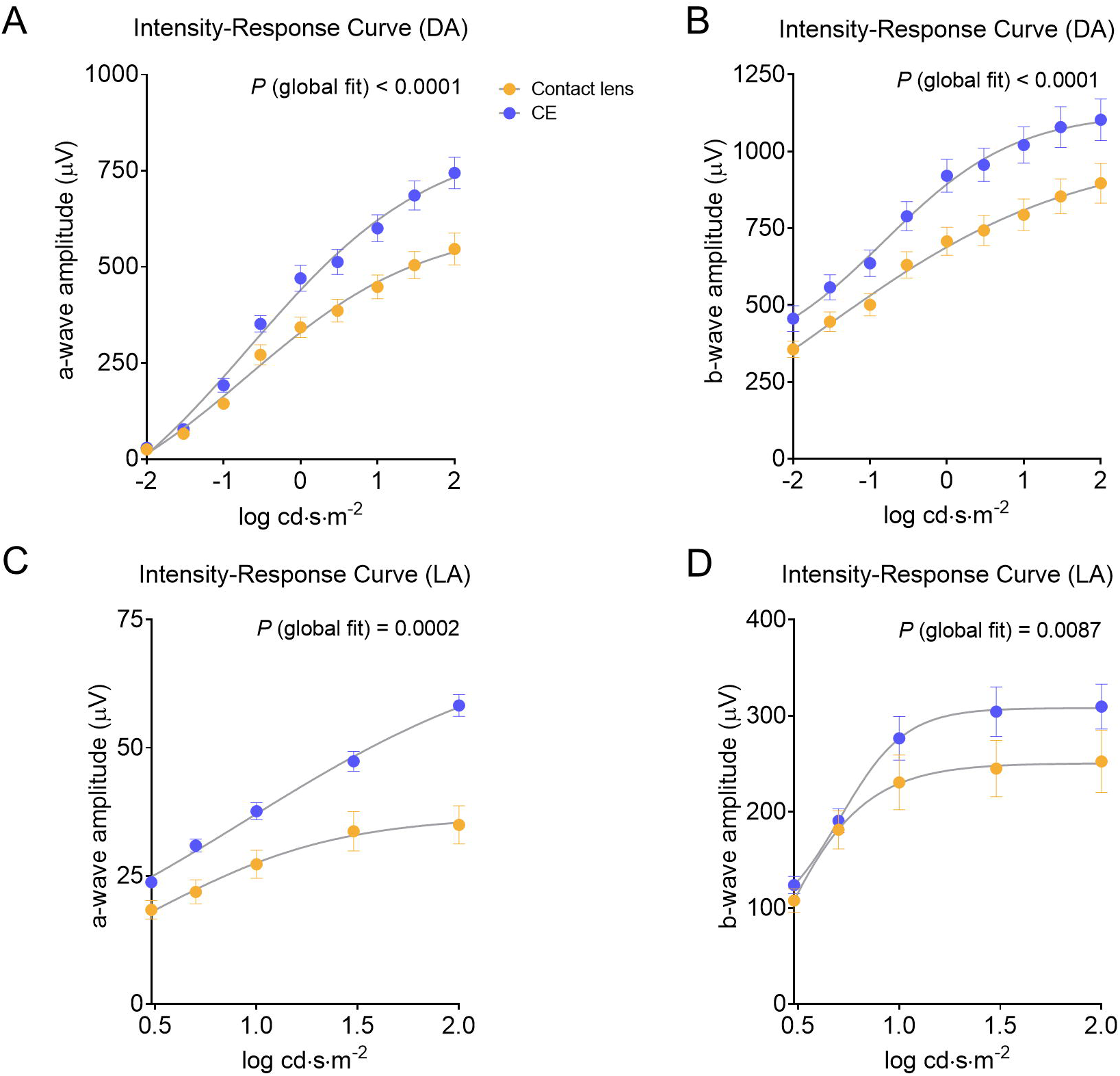
Intensity–response curves comparing a-wave and b-wave amplitudes recorded with contact lens (yellow) and CE (blue) electrodes. Intensity–response curves for a-wave (A) and b-wave (B) amplitudes under dark-adapted (DA) conditions across a flash intensity range of 0.01 to 100 cd·s·m⁻² (−2 to 2 log cd·s·m⁻²). Intensity–response curves for a-wave (C) and b-wave (D) amplitudes under light-adapted (LA) conditions across a flash intensity range of 3 to 100 cd·s·m⁻² (0.48 to 2 log cd·s·m⁻²). Datum points represent the mean ± standard error of the mean (*n* = 7). Non-linear regression curves were fitted to the data, and the statistical significance of differences between the curves was evaluated by global fitting.

### Quantitative Comparison of Scotopic Intensity–Response Curves

For quantitative comparison of dark-adapted ERG responses between the two electrodes, we analyzed the intensity–response relationships for a-wave (Figure 4A) and b-wave (Figure 4B) amplitudes as a function of flash luminance from 0.01 to 100 cd·s·m⁻² (−2 to 2 log cd·s·m⁻²). The mean amplitudes of the a-waves and b-waves increased progressively with stimulus luminance in both electrode groups. Global fitting analysis with non-linear regression was performed to determine whether the overall intensity–response functions differed significantly between the recording configurations. The global fit revealed significant difference in the intensity–response curves for both a-wave and b-wave amplitudes between the contact lens electrode and the CE electrode (*P* < 0.0001 for Figure 4A and 4B). Specifically, the mean amplitudes yielded by the CE electrode were consistently higher than those yielded by the conventional contact lens electrode across the entire luminance range tested. Similarly, under light-adapted conditions, the intensity–response curves for a-wave amplitudes were evaluated across flash intensities ranging from 3 to 100 cd·s·m⁻² (0.48 to 2 log cd·s·m⁻²). Global fitting analysis showed that the intensity–response relationships for both photopic a-wave and b-wave amplitudes differed significantly between the two electrode types (Figure 4C, a-wave: *P* = 0.0002; Figure 4D, b-wave: *P* = 0.0087). Paralleling the scotopic findings, the CE electrode recorded higher mean amplitudes than the contact lens electrode across photopic stimulus intensities. Overall, these findings demonstrated that higher ERG amplitudes were obtained with the CE electrode under both dark-adapted and light-adapted conditions.

### Individual Waveform Analysis under the DA 0.01 Condition

For further evaluation of performance of the electrodes under ISCEV standard ERG protocols, we performed the following individual paired analyses of ERG parameters at representative stimulus intensities corresponding to human clinical standards^5^: DA 0.01, DA 10, and LA 3. First, we analyzed responses elicited at a dark-adapted intensity of 0.01 cd·s·m⁻² (DA 0.01, Figure 5) to evaluate dim-flash rod-driven responses. When evaluating intensity–response relationships under dark-adapted conditions, tracking the threshold of the a-wave at low stimulus intensities (e.g., DA 0.01) provides a sensitive index for assessment of recording fidelity and signal-to-noise dynamics.^7,16,18^ Figure 5A shows the mean waveforms with shaded SEM regions recorded by both electrodes, defining the measurement criteria for a-wave and b-wave amplitudes and peak times. Paired comparisons between the contact lens and CE electrodes for individual mice revealed no statistically significant difference in a-wave amplitude (25.65 ± 2.06 μV for the contact lens electrode vs. 30.04 ± 4.02 μV for the CE electrode, *P* = 0.4688, Figure 5B), a-wave peak time (29.57 ± 0.20 ms vs. 31.64 ± 0.73 ms, *P* = 0.0625, Figure 5C), or b-wave peak time (75.29 ± 6.18 ms vs. 79.57 ± 5.50 ms, *P* = 0.1719, Figure 5E). However, the b-wave amplitude was significantly higher when recorded with the CE electrode than with the contact lens electrode (474.26 ± 37.36 μV vs. 355.95 ± 26.46 μV, *P* = 0.0156, Figure 5D). These results demonstrated that the CE electrode preserves comparable temporal response kinetics while offering signal amplitude at least equivalent to that of the conventional contact lens electrode.

**Figure 5.**
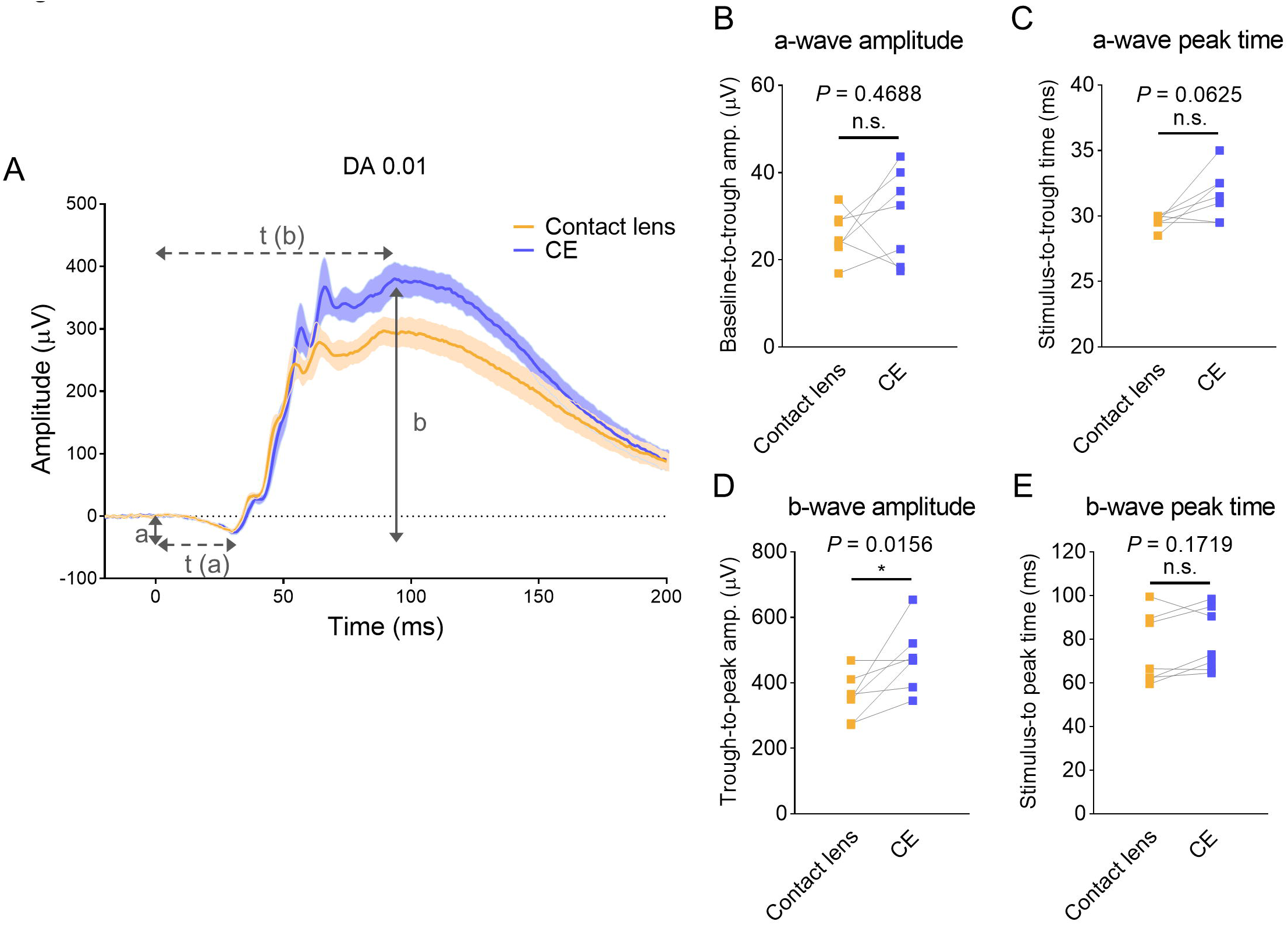
Comparison of dark-adapted (DA) electroretinographic responses recorded using contact lens and CE electrodes at a flash intensity of 0.01 cd·s·m⁻² (DA 0.01). (A) Representative grand-average electroretinographic waveforms recorded with contact lens (yellow) and CE (blue) electrodes. Shaded areas represent the mean ± standard error of the mean. Definitions for a-wave amplitude (a), b-wave amplitude (b), a-wave peak time [t(a)], and b-wave peak time [t(b)] are illustrated schematically. (B) Quantitative comparison of a-wave amplitude. (C) Quantitative comparison of a-wave peak time. (D) Quantitative comparison of b-wave amplitude. (E) Quantitative comparison of b-wave peak time. Statistical significance was determined using the Wilcoxon matched-pairs signed-rank test (*n* = 7; \**P* < 0.05; n.s., not statistically significant).

### Individual Waveform Analysis under the DA 10 Condition

Next, we examined individual paired responses at a higher dark-adapted flash intensity of 10 cd·s·m⁻² (DA 10; Figure 6), which evokes maximal combined rod and cone responses. Figure 6A shows the mean waveforms with shaded SEM regions. Paired statistical evaluations showed that the a-wave amplitude was significantly higher with the CE electrode than with the contact lens electrode (600.35 ± 35.25 μV vs. 447.99 ± 31.07 μV, *P* = 0.0313, Figure 6B). A slight but statistically significant difference was also observed in a-wave peak time (11.21 ± 0.21 ms for the contact lens electrode vs. 12.21 ± 0.15 ms for the CE electrode, *P* = 0.0469, Figure 6C), although the mean difference was minimal (approximately 1 ms) and both electrodes recorded comparable initial response kinetics. Similar to the findings for a-wave amplitude, the b-wave amplitude was significantly higher with the CE electrode than with the contact lens electrode (1020.68 ± 59.08 μV vs. 793.30 ± 51.69 μV, *P* = 0.0156, Figure 6D), whereas the b-wave peak time was not significantly different between the two configurations (32.36 ± 0.90 ms for the contact lens electrode vs. 34.36 ± 0.59 ms for the CE electrode, *P* = 0.0938, Figure 6E). Overall, these results indicated that under dark-adapted conditions with high-intensity stimuli, the response kinetics of the CE electrode are very similar to those of the contact lens electrode, while the recorded signal amplitudes tend to be larger.

**Figure 6.**
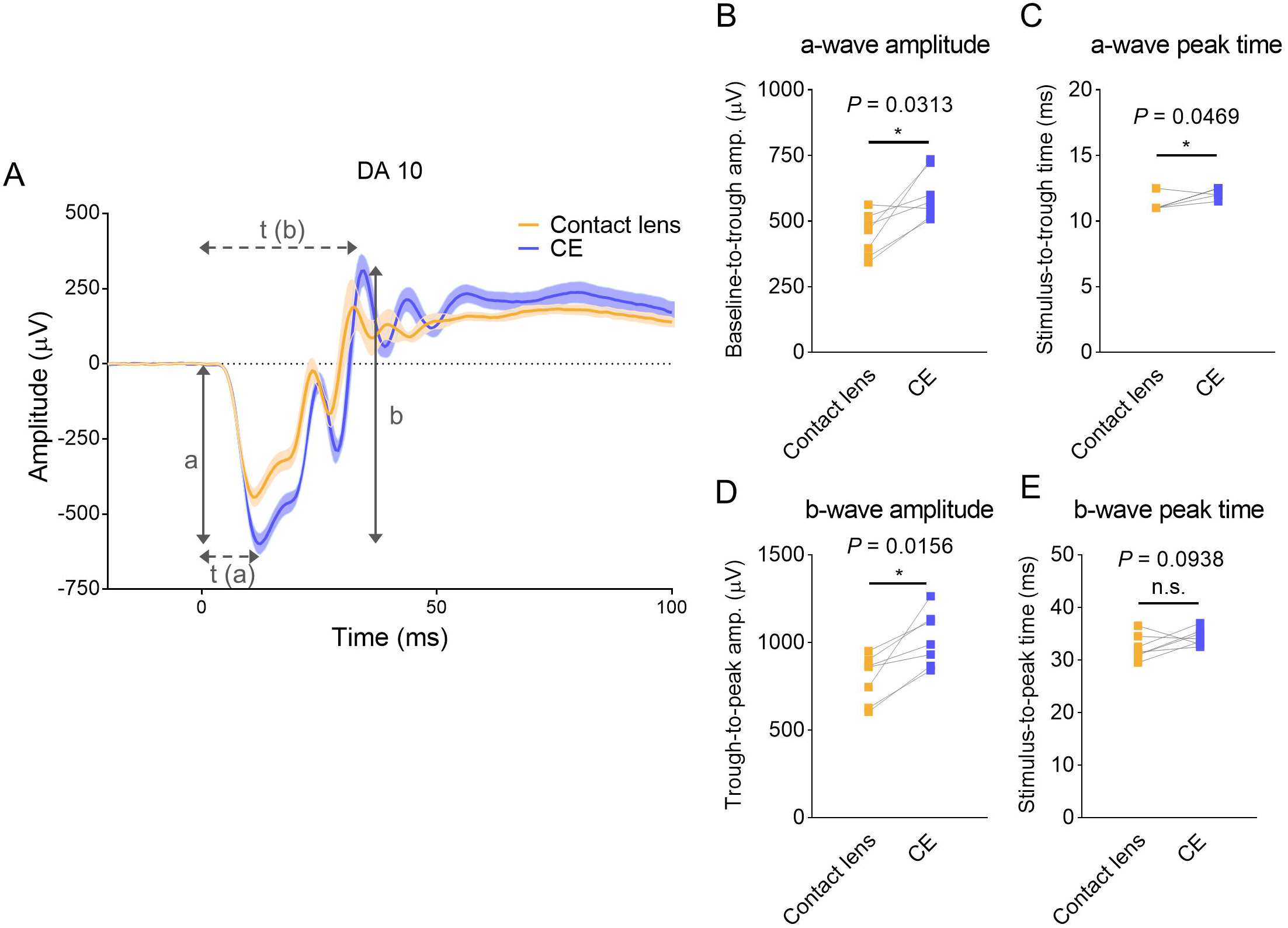
Comparison of dark-adapted (DA) electroretinographic responses recorded using contact lens and CE electrodes at a high-intensity flash of 10 cd·s·m⁻² (DA 10). (A) Representative grand-average electroretinographic waveforms recorded with contact lens (yellow) and CE (blue) electrodes. Shaded areas represent the mean ± standard error of the mean. Definitions for a-wave amplitude (a), b-wave amplitude (b), a-wave peak time [t(a)], and b-wave peak time [t(b)] are illustrated schematically. (B) Quantitative comparison of a-wave amplitude. (C) Quantitative comparison of a-wave peak time. (D) Quantitative comparison of b-wave amplitude. (E) Quantitative comparison of b-wave peak time. Statistical significance was determined using the Wilcoxon matched-pairs signed-rank test (*n* = 7; \**P* < 0.05; n.s., not statistically significant).

### Individual Waveform Analysis under the LA 3 Condition

Next, we examined individual paired responses at a light-adapted flash intensity of 3 cd·s·m⁻² (LA 3, Figure 7). Figure 7A shows the mean waveforms with shaded SEM regions. Paired statistical evaluations did not reveal any statistically significant differences between the contact lens electrode and the CE electrode for any of the parameters under the LA 3 condition, including a-wave amplitude (18.36 ± 1.83 μV vs. 22.59 ± 1.71 μV, *P* = 0.1563, Figure 7B), a-wave peak time (17.57 ± 0.51 ms vs. 18.93 ± 0.46 ms, *P* = 0.0938, Figure 7C), b-wave amplitude (107.60 ± 12.21 μV vs. 123.79 ± 9.06 μV, *P* = 0.2188, Figure 7D), and b-wave peak time (36.50 ± 2.10 ms vs. 35.36 ± 0.78 ms, *P* = 0.9688, Figure 7E). Overall, under the standard LA 3 condition, ERG responses recorded with the CE electrode were similar to those obtained with the conventional contact lens electrode.

**Figure 7.**
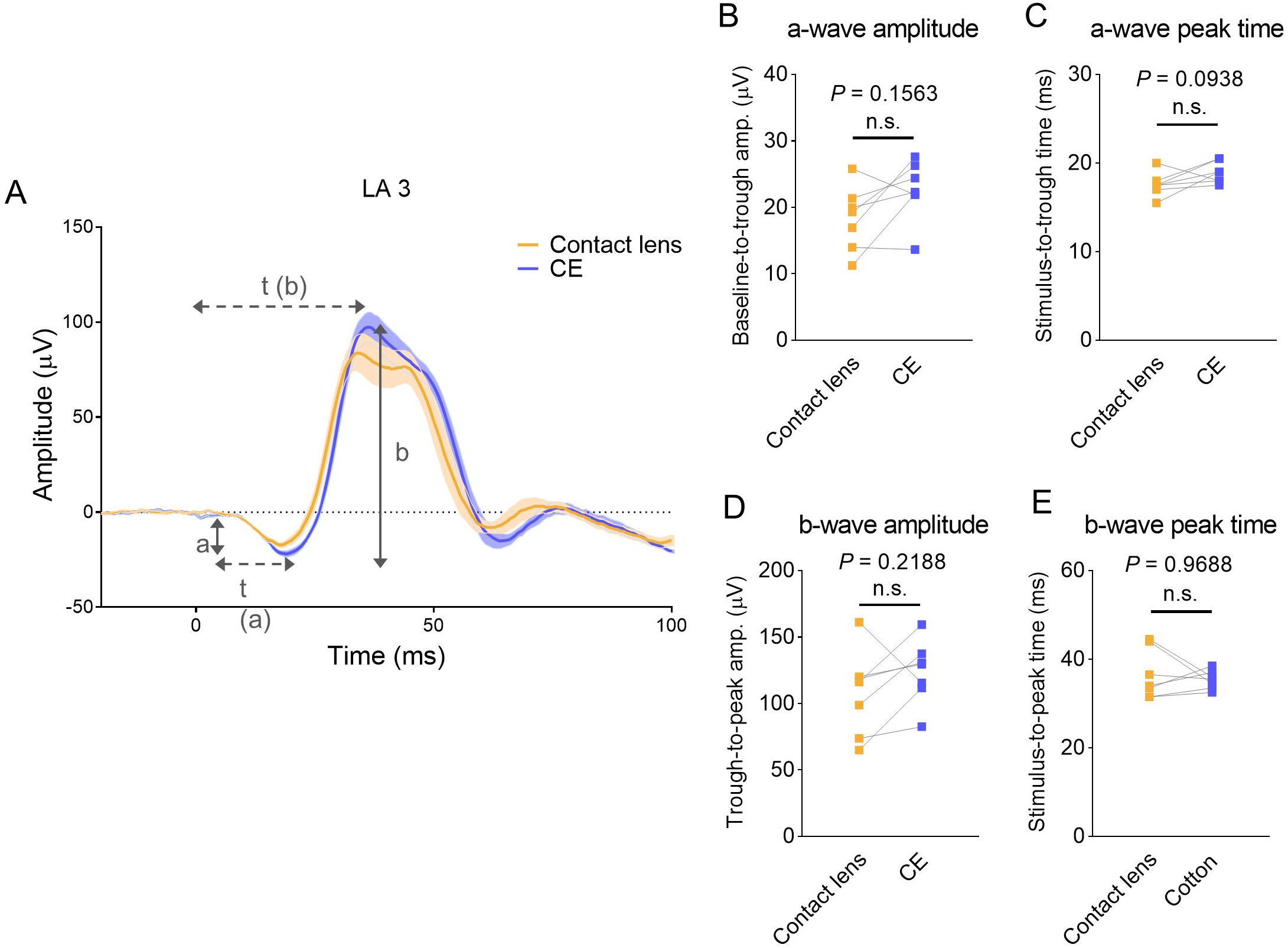
Comparison of light-adapted (LA) electroretinographic responses recorded using contact lens and CE electrodes at a flash intensity of 3 cd·s·m⁻² against a constant background light (LA 3). (A) Representative grand-average electroretinographic waveforms recorded with contact lens (yellow) and CE (blue) electrodes. Shaded areas represent the mean ± standard error of the mean. Dashed line indicates the baseline level. Definitions for a-wave amplitude (a), b-wave amplitude (b), a-wave peak time (t(a)), and b-wave peak time (t(b)) are illustrated schematically. (B) Quantitative comparison of a-wave amplitude. (C) Quantitative comparison of a-wave peak time. (D) Quantitative comparison of b-wave amplitude. (E) Quantitative comparison of b-wave peak time. Statistical significance was determined using the Wilcoxon matched-pairs signed-rank test (*n* = 7; n.s., not statistically significant).

Furthermore, while the intensity–response curve evaluated stimuli up to 100 cd·s·m⁻² to capture the full photopic dynamic range (Figure 4), we also analyzed 10 cd·s·m⁻² (LA 10) for individual paired analysis because it represents a physiologically saturated photopic stimulus that reliably evokes maximal cone-driven b-wave responses without causing excessive photobleaching (Supplementary Figure S3). Under these saturated LA 10 conditions, the two electrodes yielded generally comparable ERG responses, with the a-wave amplitude being significantly higher for the CE electrode than for the contact lens electrode (35.58 ± 2.67 μV vs. 27.25 ± 2.74 μV, *P* = 0.0313, Supplementary Figure S3B) and no significant differences in other parameters (*P* > 0.05; Supplementary Figure S3C–S3E). These results demonstrated that under saturated photopic conditions, the CE electrode provides reliable ERG responses generally comparable to those obtained with the conventional contact lens electrode, while recording a larger a-wave amplitude.

### Evaluation of Slow Ocular Response (c-wave)

ERG responses were recorded for 5 s following dark-adapted flashes across a luminance range from 0.01 to 100 cd·s·m⁻² (−2 to 2 log cd·s·m⁻²) to assess electrode performance for long-duration electrophysiological events. Following the fast a-wave and b-wave dynamics, a prominent, slow positive deflection corresponding to the c-wave, originating primarily from the retinal pigment epithelium (RPE) and Müller cells,^19–21^ was clearly observed in both contact lens and CE electrode configurations (Figure 8A and 8B; mean waveforms with shaded SEM regions shown in Supplementary Figure S4). Quantitative comparison of the c-wave intensity–response curves across the evaluated stimulus range from 0.01 to 100 cd·s·m⁻² (−2 to 2 log cd·s·m⁻²) revealed a statistically significant difference between the two electrode configurations (*P* < 0.0001, Figure 8C). While both electrodes showed comparable c-wave amplitudes at lower intensities, the mean amplitudes at moderate-to-high flash intensities were systematically higher for the CE electrode than for the contact lens electrode. These data demonstrated that the CE electrode provides stable corneal coupling comparable to that provided by the conventional contact lens electrode, enabling reliable electrophysiological recordings even during long-duration stimuli in the order of several seconds.

**Figure 8.**
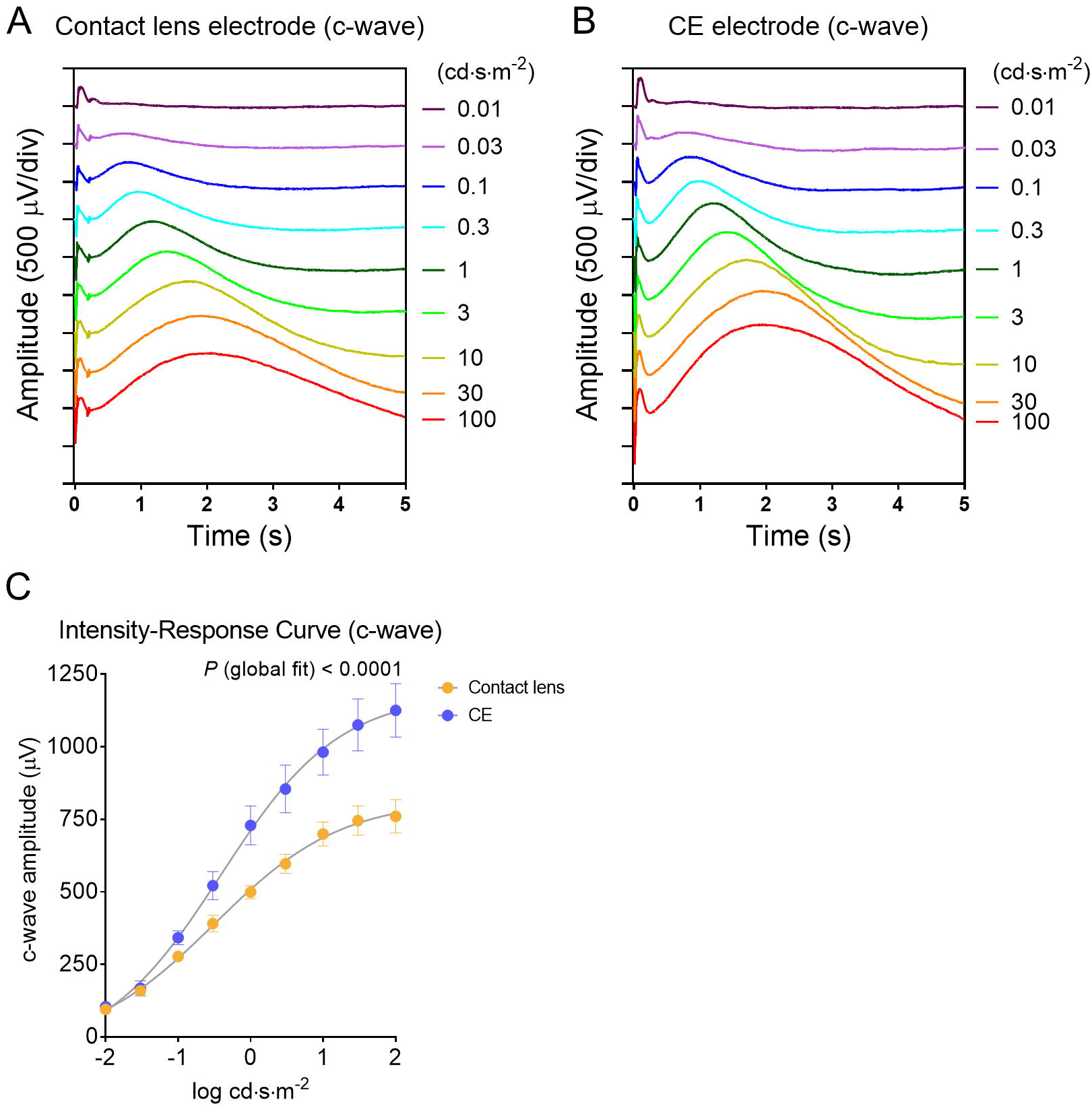
Intensity–response series of c-waves recorded using contact lens and CE electrodes across a wide range of flash intensities (0.01 to 100 cd·s·m⁻²). (A, B) Representative c-wave trace series recorded with contact lens (A) and CE (B) electrodes. Traces are stacked vertically in ascending order of flash intensity. (C) Intensity–response curves comparing c-wave amplitudes between contact lens (yellow) and CE (blue) electrodes. Datum points represent the mean ± standard error of the mean (*n* =7). Non-linear regression curves were fitted to the data, and the statistical significance of differences between the curves was evaluated using global fitting (*P* < 0.0001).

### Individual Analysis of C-Wave Waveforms under the DA 10 Condition

We performed individual paired analyses of c-wave parameters elicited by DA 10 conditions to further evaluate the response characteristics of long-duration potentials (Figure 9). Figure 9A shows the mean c-wave waveforms with shaded SEM regions over a 3.5-s recording window. Consistent with the global intensity–response relationship, paired comparisons of individual mice demonstrated that the c-wave amplitude was significantly higher when recorded using the CE electrode than when recorded using the contact lens electrode (981.21 ± 79.26 μV vs. 699.58 ± 41.53 μV, *P* = 0.0313, Figure 9B). However, there was no statistically significant difference in c-wave peak time between the contact lens and CE electrode configurations (1.66 ± 0.05 s vs. 1.73 ± 0.04 s, *P* = 0.3750, Figure 9C). These findings confirmed that while the temporal kinetics of the slow ocular response are preserved, the CE electrode offers larger amplitudes and robust signals for long-duration electrophysiological events.

**Figure 9.**
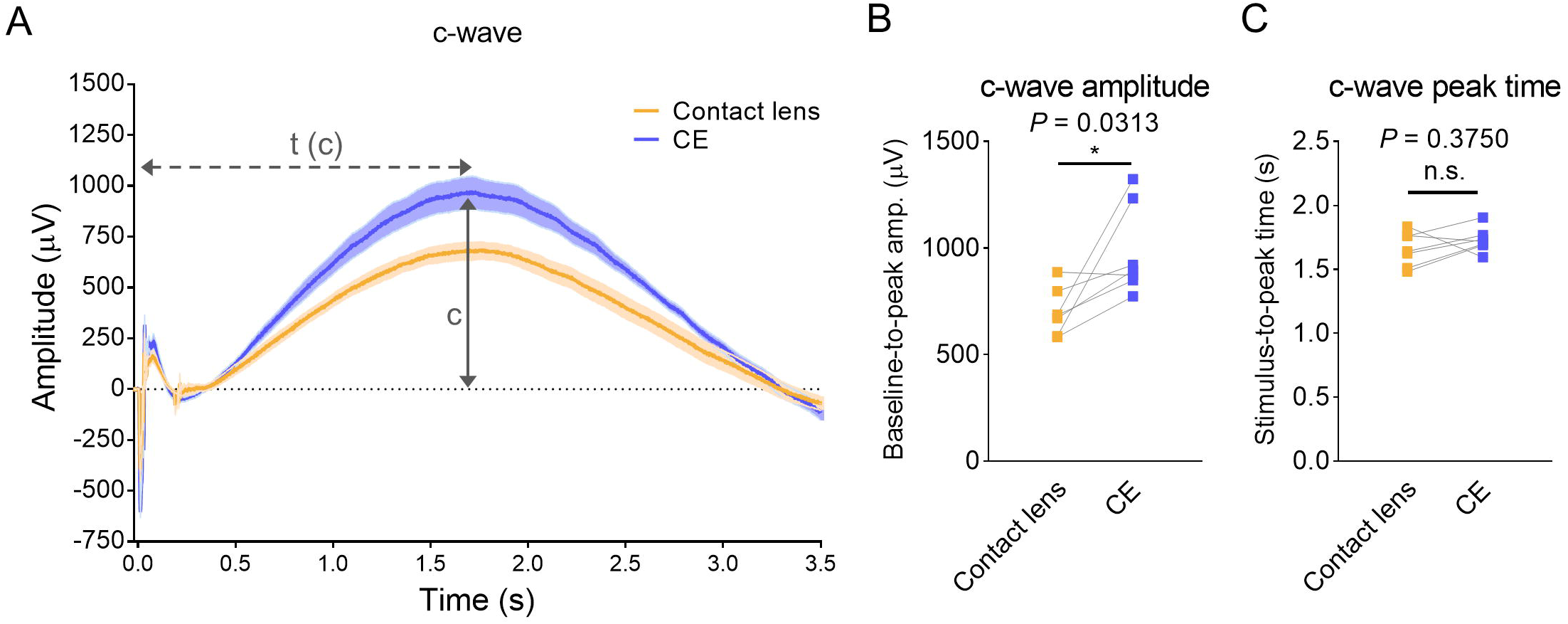
Quantitative comparison of c-wave parameters between contact lens and CE electrodes. (A) Representative grand-average c-wave waveforms recorded with contact lens (yellow) and CE (blue) electrodes. Shaded areas represent the mean ± standard error of the mean. Definitions for c-wave amplitude (c) and c-wave peak time [t(c)] are illustrated schematically. (B) Quantitative comparison of c-wave amplitude. (C) Quantitative comparison of c-wave peak time. Statistical significance was determined using the Wilcoxon matched-pairs signed-rank test (*n* = 7; \**P* < 0.05; n.s., not statistically significant).

### Evaluation of Cone-Driven Flicker ERG Responses

For evaluation of electrode performance under high-frequency repetitive stimulation, cone-driven flicker ERG responses were recorded under 20-Hz flicker stimulation, which represents the optimal frequency for isolating cone-mediated function in the rodent retina.^22^ Figure 10A shows the mean 20-Hz flicker waveforms with shaded SEM regions over a 1100-ms recording period for both the contact lens and CE electrode configurations. Quantitative analyses were conducted on the steady-state responses within the analyzed window (responses to flashes 6–15). Paired comparisons between the contact lens and CE electrode configurations revealed no statistically significant difference in trough-to-peak amplitude (68.71 ± 3.84 μV vs. 75.27 ± 2.48 μV, *P* = 0.1563, Figure 10B) or peak time (40.46 ± 0.77 ms vs. 40.54 ± 0.78 ms, *P* = 0.7813, Figure 10C). Electrode performance was also evaluated under 30-Hz flicker stimulation corresponding to the human ISCEV clinical standard. Similar to the findings for 20 Hz, the CE electrode produced 30-Hz flicker waveforms that closely overlapped with those of the contact lens electrode (Supplementary Figure S5). Quantitative analyses of steady-state responses confirmed that the response dynamics and signal amplitudes were equivalent between the two electrode systems, with no statistically significant differences in amplitude or peak time (Supplementary Figure S5B and S5C). These results demonstrated that the CE electrode accurately tracks high-frequency cone-mediated signals without introducing temporal delay or signal attenuation across both rodent-optimized (20 Hz) and clinical standard (30 Hz) flicker frequencies.

**Figure 10.**
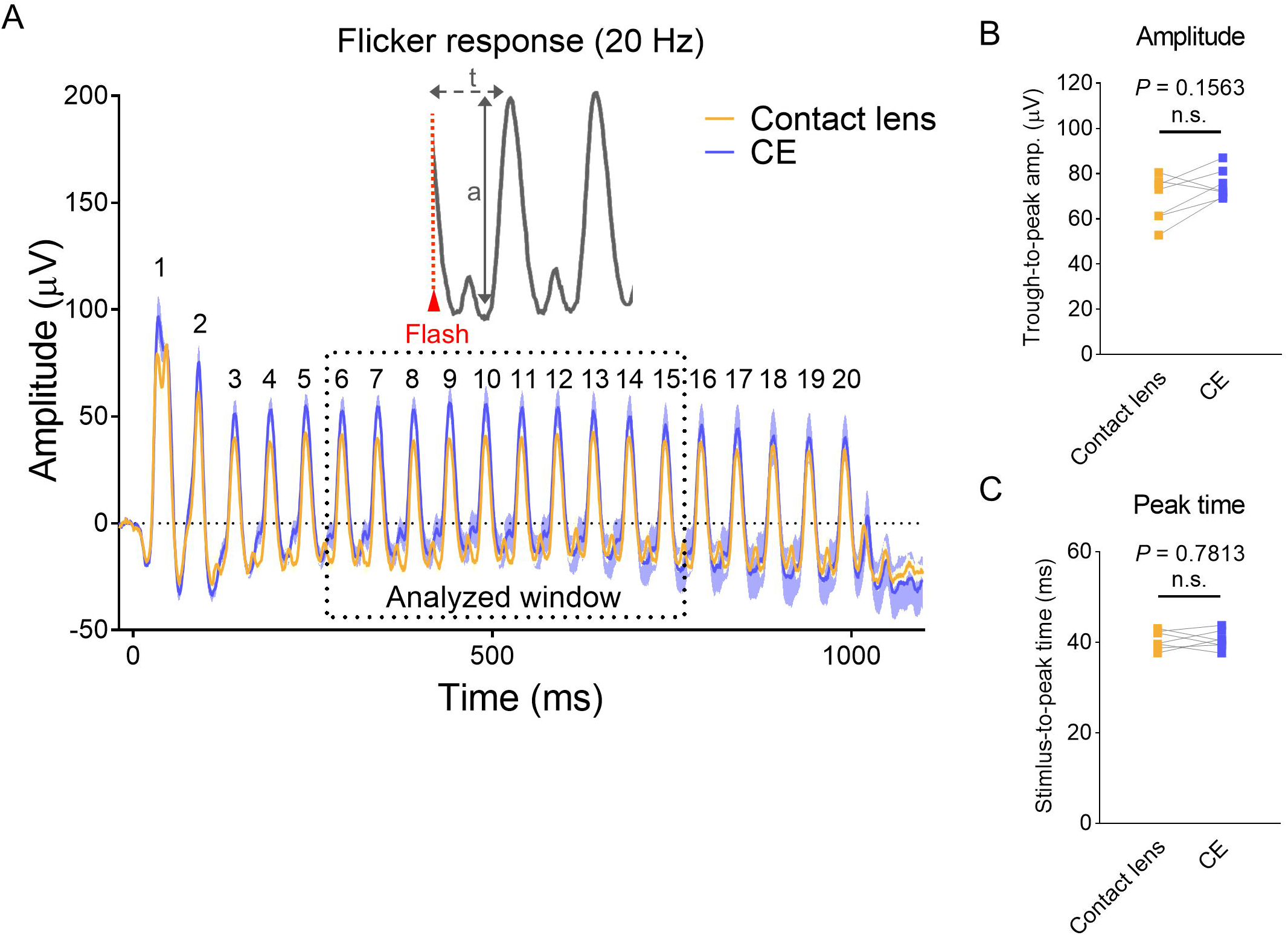
Comparison of photopic 20-Hz flicker electroretinographic responses recorded using contact lens and CE electrodes. (A) Representative grand-average flicker electroretinographic waveforms recorded with contact lens (yellow) and CE (blue) electrodes in response to 20 consecutive flash stimuli. Shaded areas represent the mean ± standard error of the mean. Responses for flashes 6 to 15 were defined as the analyzed window (dotted rectangle). The inset schematically illustrates definitions for trough-to-peak amplitude (a) and stimulus-to-peak time (t). (B) Quantitative comparison of flicker response amplitude. (C) Quantitative comparison of flicker response peak time. Statistical significance was determined using the Wilcoxon matched-pairs signed-rank test (*n* = 7; n.s., not statistically significant).

### Inter-Eye Correlation Analysis

Finally, linear regression analysis was performed on ERG amplitudes simultaneously recorded from the left and right eyes to evaluate the inter-eye recording consistency and bilateral signal reliability.^12,16^ Under dark-adapted conditions (Figure 11A–11D), the conventional contact lens electrode was found to have good inter-eye linearity for the a-wave (R^2^ = 0.8383) but showed a weaker correlation for the b-wave (R^2^ = 0.4734). In contrast, the CE electrode yielded a high inter-eye correlation across both a-wave (R^2^ = 0.9070) and b-wave (R^2^ = 0.7355) amplitudes. Similarly, under light-adapted conditions (Supplementary Figure S6A–S6D), the conventional contact lens electrode demonstrated a moderate inter-eye correlation for both photopic a-wave (R^2^ = 0.6186) and b-wave (R^2^ = 0.5245) responses, with regression slopes deviating from 1.0. In contrast, the CE electrode maintained high bilateral consistency for both photopic a-wave (R^2^ = 0.7095) and b-wave (R^2^ = 0.7236) responses. These results demonstrated that the CE electrode affords bilateral recording consistency that is comparable to or better than that of the conventional contact lens electrode under both dark-adapted and light-adapted conditions.

**Figure 11.**
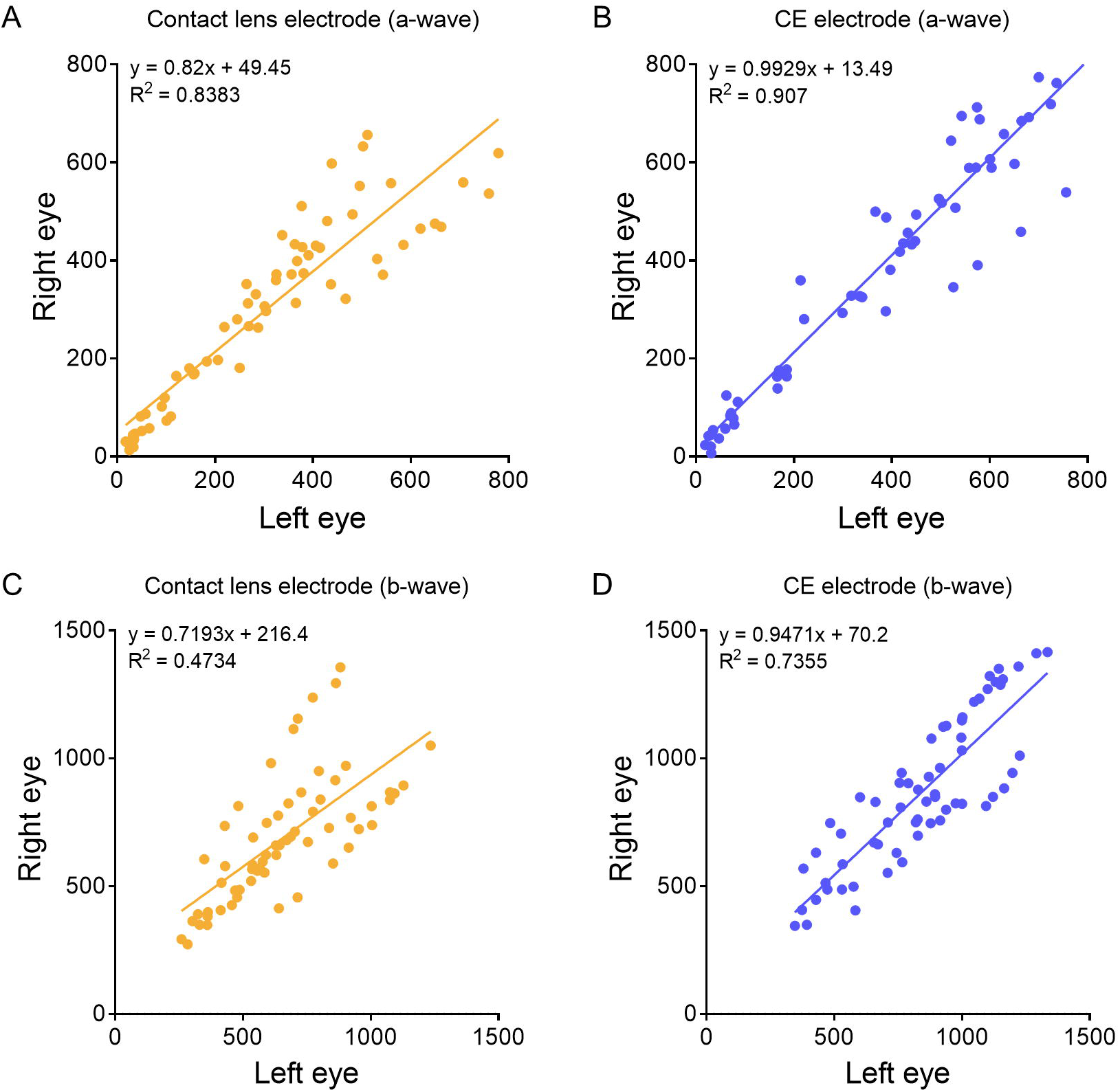
Inter-eye correlation analysis of dark-adapted electroretinographic amplitudes simultaneously recorded from contralateral eyes. Scatter plots show electroretinographic amplitudes from the left eye (x-axis, in μV) plotted against those from the right eye (y-axis, in μV) for individual mice using conventional contact lens (yellow) and CE (blue) electrodes. a-wave amplitudes recorded with contact lens (A) and CE (B) electrodes across flash intensities. b-wave amplitudes recorded with contact lens (C) and CE (D) electrodes. Solid lines represent linear regression fits. Linear regression equations and coefficient of determination (R^2^) values are indicated on each plot.

### Analysis of Mechanism of Benchtop Electrical Impedance Profiles and Inter-Electrode Consistency

We investigated the mechanisms underlying the higher overall response amplitudes (particularly the prominent c-wave) observed with CE electrodes as well as the imbalance in amplitude seen on the two sides with contact lens electrodes (Figure 11, Supplementary Figure S6) by measuring benchtop electrical impedance across a range of frequencies (Figure 12). Conventional contact lens electrodes demonstrated high impedance at low frequencies (∼20 kΩ at 1 Hz and ∼3.2 kΩ at 10 Hz) along with marked unit-to-unit variability (∼2-fold difference at 10 Hz; unit 1, ∼2.2 kΩ vs. unit 2, ∼4.1 kΩ), which decreased to 243–1,026 Ω at 1 kHz. In contrast, CE electrodes showed a consistently low, flat impedance profile from 1 Hz to 1 kHz (336–608 Ω at 10 Hz; 269–487 Ω at 1 kHz), high inter-unit consistency (with values maintained within 300– 600 Ω across frequencies), and excellent repeatability (SEM <20 Ω across 5 consecutive technical replicates).

**Figure 12.**
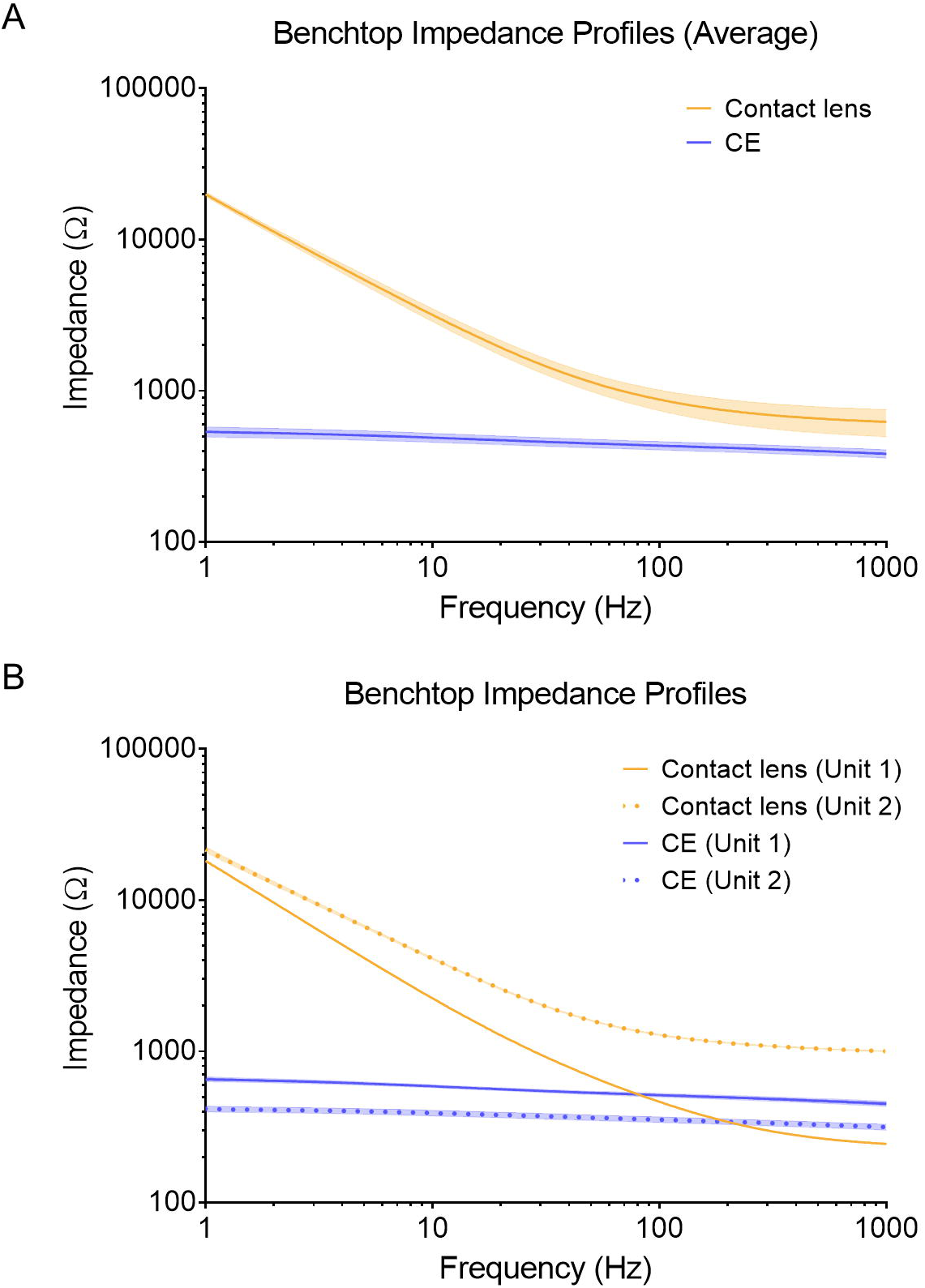
Benchtop electrochemical impedance spectra for conventional contact lens electrodes and CE electrodes. (A) Frequency-dependent impedance profiles (1 Hz to 1 kHz) measured in normal saline using an impedance analyzer (IM3590). Data represent the mean ± standard error of the mean for each type of electrode (10 measurements per group; 5 consecutive technical replicates for each of the two units). (B) Impedance profiles for each electrode unit (contact lens electrodes, units 1 and 2; CE electrodes, units 1 and 2). Each curve represents the mean ± standard error of the mean of 5 consecutive technical replicates for each unit.

## Discussion

### Baseline Noise Characteristics and Stability of the Corneal Interface

In this study, we validated a commercially available auditory canal (CE) electrode, originally designed for human electrocochleography, as a practical and highly user-friendly alternative to conventional corneal contact lens electrodes for mouse ERG. It is important to establish a stable electrical baseline at the corneal interface for accurate signal quantification before evaluating light-evoked retinal responses. As shown in Figure 2C, there was no statistically significant difference in the root mean square noise level between the contact lens electrode and the CE electrode. Both electrode configurations demonstrated high electrical background stability. Notably, the baseline noise levels recorded were fully comparable to the fundamental noise floor (approximately 20 μV) reported in functionally unresponsive mouse models under standard testing conditions.^23^ For strict isolation of intrinsic biophysical properties and avoidance of the signal variability associated with viscous coating medium, high-viscosity lubricants (e.g., methylcellulose or hydroxypropyl methylcellulose) were intentionally omitted, and both electrodes were wet exclusively with 0.9% saline. Under these standardized conditions, the non-woven fabric pad maintained a stable conductive liquid layer across the cornea, eliminating displacement artifacts and providing a high signal-to-noise ratio without requiring viscous agents.

### Functional Comparability and Intensity-Dependent Dynamics Across Recording Platforms

A prominent characteristic of the CE electrode system is its clear demonstration of light intensity-dependent responses under both dark-adapted and light-adapted conditions (Figures 3 and 4). Across a broad range of flash intensities, both the contact lens and CE electrode systems generated smooth, well-characterized non-linear intensity–response curves for dark-adapted and light-adapted a-waves and b-waves (Figure 4A–4D). Importantly, the light-evoked responses recorded with the CE electrode showed robust stimulus–intensity responsiveness similar to not only the conventional contact lens electrode used in this study but also to previously reported contact lens systems,^6,9^ gold wire loop electrodes,^7,18^ and dedicated testing platforms such as the Celeris system.^10,13^

### High Signal Detection Efficiency Under Standard Dark-Adapted and Light-Adapted ERG Protocols

We systematically assessed ERG responses under dark-adapted (DA 0.01 and DA 10) and light-adapted (LA 3) conditions to evaluate the physiological comparability of the two electrode systems based on human clinical ISCEV standards (Figures 5–7). Overall, the CE electrode was found to have functional performance that was fully comparable, and in terms of signal amplitude, superior to that of the conventional contact lens electrode. Under dark-adapted conditions, the dim-flash rod response (DA 0.01) showed overall waveforms and amplitudes that were very similar between the two electrode systems, confirming functional equivalence in evaluation of rod pathway sensitivity (Figure 5). For the maximum flash response (DA 10), the CE electrode yielded a-wave and b-wave amplitudes that were higher than those yielded by the contact lens electrode (Figure 6B and 6D). A minor statistically significant difference in a-wave peak time (approximately 1 ms) was observed (Figure 6C); however, this difference fell well within the normal range reported by previous ERG studies in mice^9,13,16,18,24^ and does not reflect a meaningful physiological difference or delay in photoreceptor response kinetics.

Under light-adapted conditions (LA 3), the two electrode systems captured virtually equivalent photopic responses (Figure 7). Furthermore, testing under a saturated photopic stimulus (LA 10) produced generally comparable waveforms, with the CE electrode recording a slightly higher a-wave without compromising overall response kinetics (Supplementary Figure S3). Together, these findings demonstrated that the CE electrode reliably delivers high-quality ERG recordings that are fully equivalent or superior to the conventional contact lens electrode across all standard clinical stimulus conditions.

### Biophysical Mechanisms and Inter-Study Variations in ERG Signal Amplitudes

The enhanced rod and cone response amplitudes observed with the CE electrode in this study align well with previous literature demonstrating that ERG signal amplitude depends on the specific electrode configuration and materials used.^12,16^ In 2011, Chekroud et al. reported that cotton-wick electrodes recorded consistently higher a-wave and b-wave amplitudes than metallic wire loop electrodes, including gold, silver, and copper.^16^ Furthermore, conventional corneal contact lens electrodes frequently incorporate gold-based conducting elements, which are characterized by significantly higher impedance and more capacitive behavior than silver-based or fabric-based electrodes, making them more susceptible to signal attenuation.^11^ The lower interface impedance and stable capillary hydration of the CE electrode likely account for its enhanced signal pickup efficiency. Indeed, our benchtop impedance analysis confirmed that the impedance of the CE electrode was lower than that of the conventional contact lens electrode in our experimental setup, which is consistent with the findings of Man et al.¹¹

This heightened detection sensitivity is also prominent in the a-wave parameters. For example, in 2023, under maximum DA 10 conditions, Lindovsky et al. reported a mean a-wave amplitude of 176.3 ± 62.3 μV in 10-week-old C57BL/6 mice using gold ring electrodes.^18^ In contrast, our setup yielded a mean a-wave amplitude of 447.99 ± 82.21 μV with the conventional contact lens electrode and 600.35 ± 93.27 μV with the CE electrode (expressed as the mean ± standard deviation for direct comparison with the findings of Lindovsky et al.), while maintaining a-wave peak times (approximately 11–13 ms) that remain similar to historical values (approximately 14–15 ms). Absolute ERG amplitudes and peak times often vary across studies as a result of differences in mouse strains used,^23^ age,^10^ type of anesthesia,^24^ and specific recording environments.^13^ Nevertheless, because both types of electrode were evaluated in the same animals under identical recording conditions in a paired design, the significantly larger a-wave and b-wave amplitudes captured by the CE electrode confirm its intrinsic advantage in terms of stable biophysical contact and electrical sensitivity, without compromise of the initial response kinetics.

### Enhanced Signal Pickup Sensitivity in Long-Duration c-Wave Recordings by the CE Electrode

The c-wave is a corneal-positive deflection observed following the classic a-wave and b-wave, representing the net sum of multiple cellular events, primarily the light-induced hyperpolarization of the apical membrane of the RPE and the Müller cell-derived slow PIII component.^25,26^ Under extended illumination, the c-wave is followed by a sequence of RPE-derived slow potentials, including the fast oscillation, light peak, and OFF-response.^25,27^ Capturing this entire sequence of RPE-related c-wave events requires sustained exposure to light over several (e.g., 7) minutes.^28^ However, such prolonged light exposure is not strictly necessary for specific evaluation of the c-wave alone; reliable c-wave recordings can be achieved using light stimuli lasting only a few (e.g., 5) seconds^8,29^ or brief millisecond-order flash stimuli (e.g., 20 ms)^20^ when appropriate low-frequency filter limits (e.g., 0.01–0.03 Hz) are applied. In this study, we successfully performed a simplified analysis of the c-wave elicited by a brief 1-ms flash stimulus under dark-adapted conditions (Figures 8 and 9). Although direct-coupled ERG protocols or lower high-pass filter cutoffs (0.01–0.03 Hz) have traditionally been recommended to prevent signal attenuation of slow RPE-derived potentials,^28–30^ amplifier bandpass filters in our setup were set to the hardware lower limit of 0.1 Hz because of system specifications (BioAmp). Despite this cutoff limit, robust and reproducible c-wave profiles were consistently captured across stimulus intensities (Figure 8). Evaluation of electrophysiological potentials over a time frame of several seconds to minutes requires low noise levels and minimal baseline drift. As described above, several studies have compared the performance of homemade cotton electrodes with that of various metallic electrodes, such as gold, silver, and copper wires, primarily for recording conventional flash ERG components.^12,14,16^ However, these evaluations did not validate the baseline stability of cotton-wick electrodes during slow potential measurements over a time frame of several seconds, such as the c-wave. In this study, we demonstrated that the commercially available CE electrode setup maintains an excellent signal-to-noise ratio with minimal baseline drift even during second-scale recording durations, demonstrating performance comparable or superior to that of conventional contact lens electrodes and confirming its stability and practical reliability for analysis of c-waves.

### High Compatibility of Photopic Flicker ERG Recordings

Photopic flicker ERGs are widely used in rodents as well as in humans to isolate and evaluate cone-driven functioning in the retinal pathway under repetitive light stimulation by suppressing rod-mediated responses.^10,31^ In human clinical ERG, a 30-Hz flicker stimulus has been established as the standard protocol for assessing the responsiveness of the cone system.^5^ However, in rodents, a 20-Hz flicker is also often used to obtain clear cone responses with high amplitude and minimal signal loss.^22,31^ High-frequency flicker ERGs serve as a robust index for evaluation of electrode stability because they require steady signal tracking to record each peak accurately without delay or dampening. In this study, we evaluated cone-driven function using both 20-Hz and 30-Hz photopic flicker ERGs. At both frequencies, the CE electrode produced flicker waveforms that closely matched those recorded by the conventional contact lens electrode throughout the stimulation period (Figure 10, Supplementary Figure S5). Statistical analysis did not reveal any significant difference in trough-to-peak amplitude or stimulus-to-peak time between the two electrodes. These results indicated that the CE electrode maintains stable corneal contact with accurate response timing, offering performance fully comparable to that of the contact lens electrode for evaluation of functioning in the cone pathway.

### Inter-Eye Differences Introduced by Electrode Limitations

The high inter-eye correlation coefficients reported by Chekroud et al. using cotton-wick electrodes (a-wave, R^2^ = 0.929; b-wave, R^2^ = 0.967) and gold electrodes (a-wave, R^2^ = 0.681; b-wave, R^2^ = 0.765) were derived within a relatively narrow amplitude range (a-wave, 0–200 μV; b-wave, 0–600 μV).^16^ However, our dataset spanned a wider range of response amplitudes (a-wave, 0–800 μV; b-wave, 0–1500 μV), including high-intensity responses, where there is an inherent increase in absolute amplitude variations (Figure 11). Within comparable lower amplitude ranges (e.g., 0–200 μV for the a-wave), the CE electrode similarly showed tight data clustering and high bilateral linearity. Notably, the conventional contact lens electrode showed mild inter-eye variability in signal amplitude. Considering that such bilateral discrepancies were not observed with the CE electrode under identical recording conditions, this inter-eye difference is likely electrode-derived and attributable to subtle physical variations in handmade rigid electrodes, minor shifts in corneal contact angle, or subtle differences in the contact pressure applied. While averaging responses from both eyes effectively cancels this bias in routine single-animal assessments, it introduces unwanted variation when performing direct inter-eye comparisons in interventional or unilateral disease models. These findings highlight an important practical consideration, namely, that custom-fabricated rigid electrodes may show inherent variability when used for inter-eye measurements. Acknowledging these electrode-specific limitations is essential when designing contralateral-controlled studies. In this regard, the CE electrode, which features a simple construction of silver wire wrapped around non-woven fabric, reduces variability between electrode batches and offers a reliable platform that avoids such systematic bilateral discrepancies.

Consistent with these observations, benchtop impedance analysis revealed a two-fold difference between the contact lens electrode pair (unit 1, ∼2.2 kΩ vs. unit 2, ∼4.1 kΩ at 10 Hz; Figure 12). In our bilateral recording setup, application of the lower-impedance electrode (unit 1) to the right eye coincided with signal amplitudes that were larger than those obtained with the higher-impedance electrode (unit 2) on the left eye, which likely contributed to the shift in the y-intercept and reduced bilateral linearity (R^2^). Furthermore, the consistently low impedance of the CE electrodes (down to 1 Hz) may have facilitated low-frequency signal retention, offering a plausible mechanism for their c-wave amplitudes being larger than those of the contact lens electrodes. Overall, these findings suggest that the impedance characteristics are a key factor influencing signal magnitude and symmetry in bilateral ERG assessments.

### An Easy-to-Use Non-Woven Fabric Electrode for Mouse ERG

Another key advantage of the CE electrode is its ease of handling and versatility. When using conventional contact lens electrodes, precise positioning and careful control of corneal pressure are essential because excessive pressure on the eye can alter intraocular pressure and reduce ERG amplitudes.^8,15^ Fluid-filled chamber systems were developed to provide a non-contact wet electrode for clinical diagnostic applications, thereby minimizing corneal invasiveness and mechanical contact^32^; however, such systems are structurally complex, vulnerable to leakage of liquid, and impractical for small rodent eyes. Furthermore, contact lens electrodes require strict selection of size to match growth of the eye; changing electrode sizes during longitudinal studies can alter corneal fit and introduce variability into recorded waveforms.^8^ In contrast, the CE electrode naturally conforms to a wide range of eye sizes without exerting excessive pressure or requiring precise size matching. By retaining saline, the CE electrode smoothly conforms to the corneal surface, minimizing mechanical contact. Consequently, this design provides a practical balance, offering signal fidelity similar to that of conventional contact lens electrodes while maintaining the corneal safety associated with fluid-filled systems. Importantly, even when re-application or readjustment of an electrode is required in the darkroom, the setup remains straightforward: the CE electrode can be readily placed after moistening the cornea with saline, followed by gently absorbing excess fluid. Indeed, our parallel study confirmed that this CE electrode enabled reliable recording of scotopic responses (e.g., DA 0.01 and DA 10) even in young mice shortly after weaning (around postnatal day 20). The same electrode configuration can be applied seamlessly from early development through to advanced age without requiring size matching, providing a highly consistent interface for longitudinal studies, such as tracking normal aging or progression of chronic disease.^9,10^ Furthermore, this commercially available electrode contributes substantially to standardization of data and high inter-laboratory reproducibility because of its standard structure.

Although originally designed for single-use in clinical diagnostic applications, this CE electrode can be readily re-sterilized and re-used multiple times in animal research; notably, unlike conventional contact lens electrodes, it can be readily replaced whenever its electrical properties start to deviate from the initial baseline. However, no such replacement was necessary throughout this study: the same set of electrodes was used repeatedly from initial preliminary testing through to final data acquisition, during which no signal degradation, waveform instability, or abnormal ERG amplitudes or peak times were observed. This combination of structural consistency, ease of handling, and practical durability renders the CE electrode a highly accessible and standardized tool for routine visual electrophysiology.

## Conclusion

The commercially available CE electrode serves as a practical, cost-effective, and reliable alternative to the conventional contact lens electrode for murine ERG. It achieves functional equivalence in waveform fidelity and response kinetics across standard scotopic, photopic, and high-frequency flicker conditions. Notably, the CE electrode maintains a stable corneal interface with minimal baseline drift, ensuring accurate recording of long-duration potentials such as the c-wave. Furthermore, the CE electrode minimizes systematic inter-eye discrepancies, showing high bilateral correlation suitable for contralateral-controlled studies. Moreover, its consistent manufacturing quality and reusability following sterilization overcome the structural variability of traditional handmade cotton wicks, thereby supporting data standardization and promoting inter-laboratory reproducibility. This practical system serves as a robust tool in vision research laboratories, improving both experimental efficiency and data consistency.

## Acknowledgments

This work was supported by a Grant-in-Aid for Scientific Research from the Japan Society for the Promotion of Science (JSPS KAKENHI grant number JP25K12901 to Y.I.), a research project grant (R07B091) from Kawasaki Medical School to Y.I., and the Japan Agency for Medical Research and Development (AMED) under grant number JP26ym0126810 to J.M. We thank Mitsuhiro Abe (Unique Medical) and Shuhei Mashimo (Bio Research Center) for their technical support in setting up the experimental system. We also thank Edanz (https://jp.edanz.com/ac) for editing a draft of this manuscript.

## Supplementary information

**Supplementary Figure S1.**
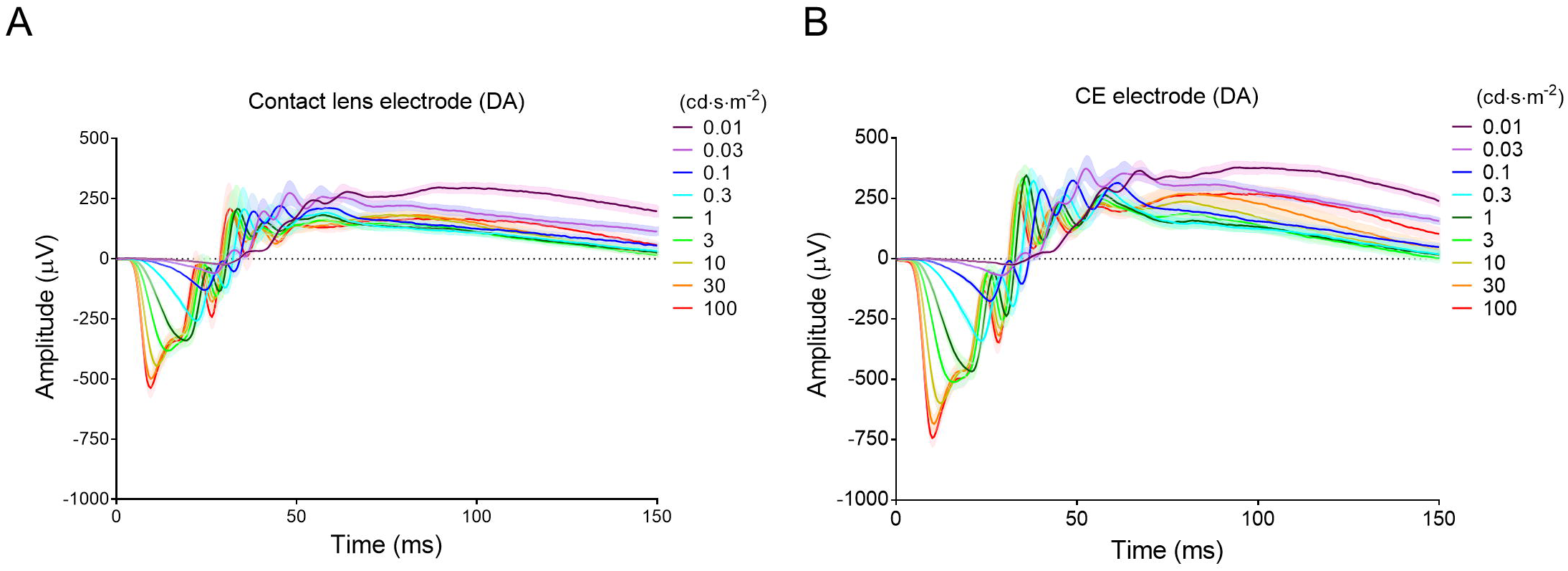
Dark-adapted (DA) electroretinographic response series across various flash intensities recorded using contact lens and CE electrodes. (A, B) Grand-average electroretinographic waveforms recorded with contact lens (A) and CE (B) electrodes across a flash intensity range of 0.01 to 100 cd·s·m^−2^ under DA conditions. Waveforms corresponding to various flash intensities are color-coded according to the inset legend. Shaded areas represent the mean ± standard error of the mean. (*n* = 7).

**Supplementary Figure S2.**
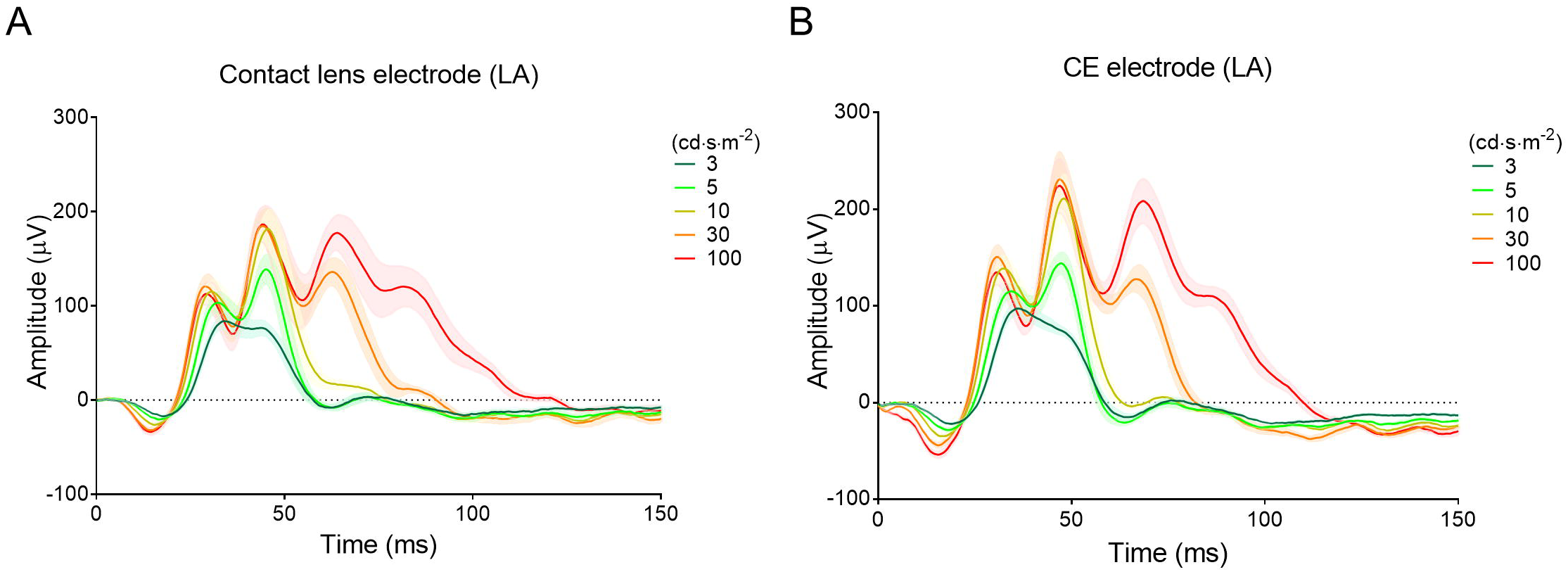
Light-adapted (LA) electroretinographic response series across various flash intensities recorded using contact lens and CE electrodes. Grand-average electroretinographic waveforms recorded with contact lens (A) and CE (B) electrodes across a flash intensity range of 3 to 100 cd·s·m^−2^ on a constant background light under LA conditions. Waveforms corresponding to different flash intensities are color-coded according to the inset legend. Shaded areas represent the mean ± standard error of the mean (*n* = 7).

**Supplementary Figure S3.**
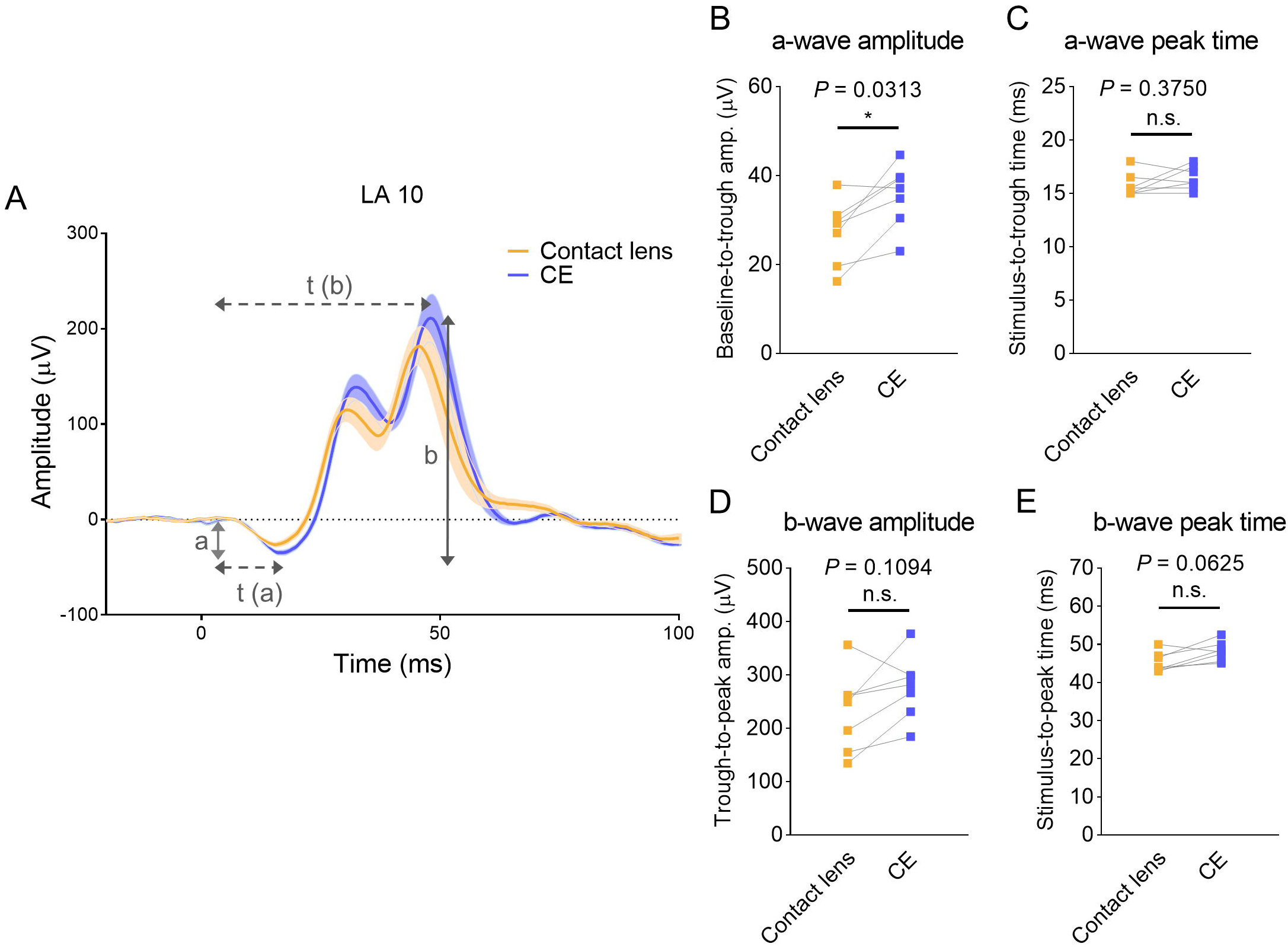
Comparison of light-adapted (LA) electroretinographic responses recorded using contact lens and CE electrodes at a high-intensity flash of 10 cd·s·m^−2^ against a constant background light (LA 10). (A) Representative grand-average electroretinographic waveforms recorded with contact lens (yellow) and CE (blue) electrodes. Shaded areas represent the mean ± standard error of the mean. Definitions for a-wave amplitude (a), b-wave amplitude (b), a-wave peak time [t(a)], and b-wave peak time [t(b)] are schematically illustrated. (B) Quantitative comparison of a-wave amplitude. (C) Quantitative comparison of a-wave peak time. (D) Quantitative comparison of b-wave amplitude. (E) Quantitative comparison of b-wave peak time. Statistical significance was determined using the Wilcoxon matched-pairs signed-rank test (*n* = 7; \**P* < 0.05; n.s., not statistically significant).

**Supplementary Figure S4.**
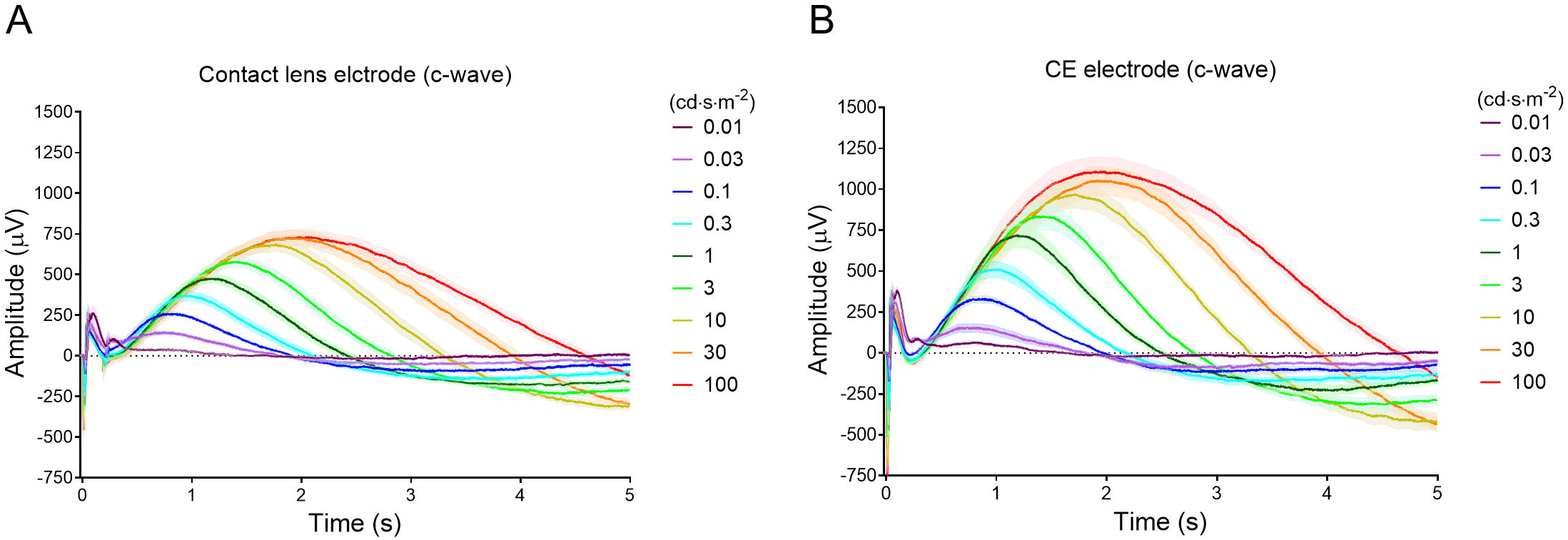
Dark-adapted c-wave response series across various flash intensities recorded using contact lens and CE electrodes. Grand-average c-wave waveforms recorded with contact lens (A) and CE (B) electrodes across a flash intensity range of 0.01 to 100 cd·s·m^−2^ under DA conditions. Waveforms corresponding to various flash intensities are color-coded according to the inset legend. Shaded areas represent the mean ± standard error of the mean (*n* = 7).

**Supplementary Figure S5.**
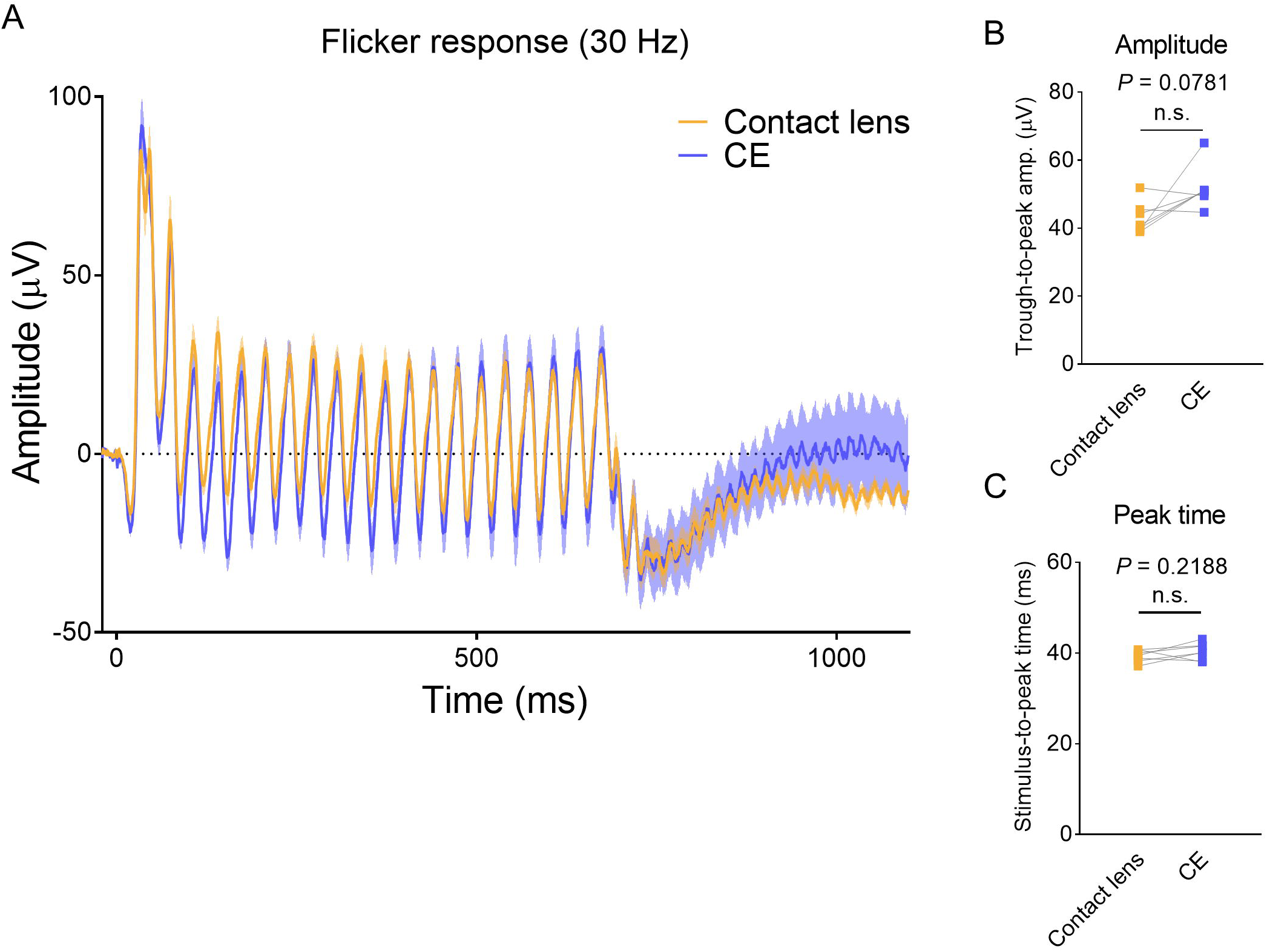
Comparison of photopic 30-Hz flicker electroretinographic responses between contact lens and CE electrodes. (A) Representative grand-average flicker electroretinographic waveforms recorded with contact lens (yellow) and CE (blue) electrodes. Shaded areas represent the mean ± standard error of the mean. (B) Quantitative comparison of flicker response amplitude. (C) Quantitative comparison of flicker response peak time. Statistical significance was determined using the Wilcoxon matched-pairs signed-rank test (*n* = 7; n.s., not statistically significant).

**Supplementary Figure S6.**
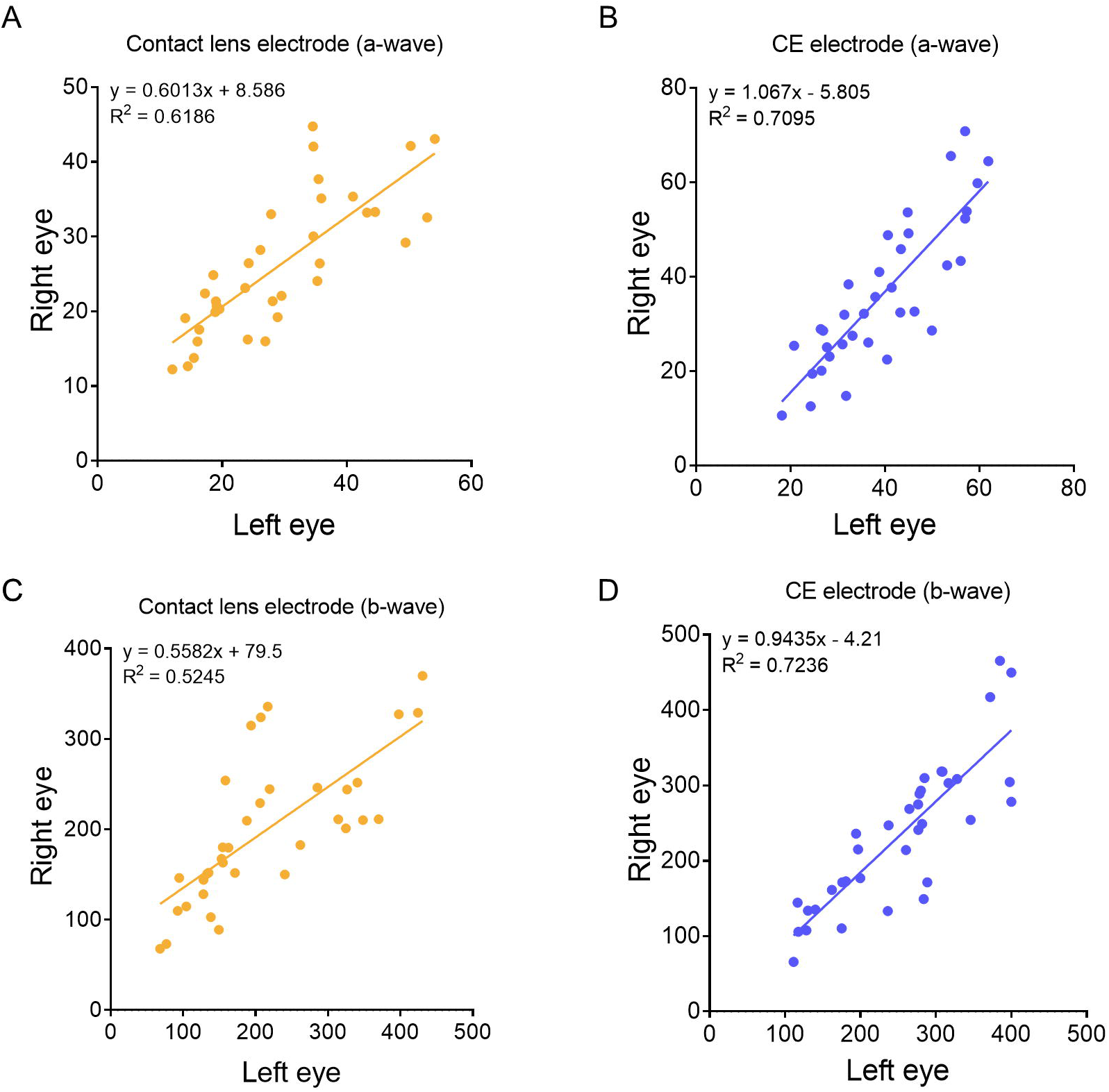
Inter-eye correlation analysis of light-adapted electroretinographic amplitudes simultaneously recorded from contralateral eyes. Scatter plots show electroretinographic amplitudes from the left eye (x-axis, in μV) plotted against those from the right eye (y-axis, in μV) for individual mice (*n* = 7) using conventional contact lens (yellow) and CE (blue) electrodes. a-wave amplitudes recorded with contact lens (A) and CE (B) electrodes across flash intensities. b-wave amplitudes recorded with contact lens (C) and CE (D) electrodes. Solid lines represent linear regression fits. Linear regression equations and coefficient of determination (R^2^) values are indicated on each plot.

